# Individual differences in artificial neural networks capture individual differences in human behavior

**DOI:** 10.64898/2026.02.10.705061

**Authors:** Herrick Fung, N. Apurva Ratan Murty, Dobromir Rahnev

**Author notes:** **Correspondence:** Herrick Fung, School of Psychological and Brain Science Georgia Institute of Technology, 654 Cherry Str NW Atlanta, GA 30332.

## Abstract

Human behavior differs substantially across individuals. While artificial neural networks (ANNs) are regarded as promising models of human perception, they are often assumed to lack such individual differences. Here, we demonstrate that multiple instances of the same ANN architecture exhibit substantial individual differences in behavior that mimic those observed in humans. We trained and tested 60 ANN instances from three architectures on a digit recognition task and found notable individual differences in overall accuracy, confidence, and response time (RT). Critically, these individual differences in ANN instances mapped consistently onto the individual differences produced by 60 humans performing the same task, with the mapping strength often approaching the human-to-human benchmark across all three behavioral metrics (accuracy, confidence, RT). The mapping generalized even across behavioral metrics: an ANN instance that aligned with an individual human on accuracy also aligned with the same individual on confidence and RT. These findings extended to a more complex, 10- choice blurry object recognition task, though the human-ANN mapping was generally less robust than the human-human benchmark. Overall, these findings open the possibility of using ANN ensembles as computational proxies for probing the mechanisms underlying human variability.

## 1 Introduction

No two individuals perceive, think, or act in the same way. Understanding and predicting individual differences in human behavior is a critical challenge in the quest to uncover the complexity of human intelligence. In recent years, artificial neural networks (ANNs) have emerged as the leading computational models of the primate visual system and perceptual decision making for complex images (Kriegeskorte, 2015; McGrath et al., 2024; Wichmann & Geirhos, 2023). Yet, since Turing’s original proposal of the imitation game (Turing, 1950), AI and, more recently, NeuroAI (Feather et al., 2025) have largely aimed to model the behavior of an *average* human. Whether ANNs can capture human behavior at the individual level, that is, the full spectrum of human behavioral diversity, is yet to be explored.

Indeed, ANNs are often considered uniform within a given architecture (e.g., AlexNet), with little meaningful variability between different model instances. It is only recently that studies have challenged this assumption, demonstrating that even within a fixed ANN architecture, different random weight initializations (Mehrer et al., 2020), different orderings of training images, and different frequencies of items and categories (Chow & Palmeri, 2024) can lead to substantial differences in the internal representations. Critically, these representational differences are likely to translate into behavioral variability and several recent papers have found clear behavioral differences between ANN instances that only differ in their random initializations (Baek et al., 2021; Green et al., 2026; Rafiei et al., 2024; Shekhar & Rahnev, 2024).

While differences between ANN instances are beginning to gain attention, it remains unclear whether such individual differences in ANNs can be meaningfully related to individual differences in human behavior. Although some studies have argued for the potential relevance of individual differences across ANNs to individual differences in human behavior (Annis et al., 2021; Chow & Palmeri, 2024), no studies to date have directly compared individual differences in humans and ANNs. If individual differences in ANNs mimic those observed in humans, these networks could potentially serve as proxy models for human variability, supporting advances in both theoretical insight and empirical investigation. For instance, by manipulating model instances and testing their alignment with individuals, we could use ANNs to reverse engineer the origins of individual differences in humans, offering a new approach to probe the mechanisms underlying behavioral variability (Annis et al., 2021; Chow & Palmeri, 2024; Strock et al., 2025; Wood et al., 2024). A proxy model for human variability would also allow new *in silico* experiments to precisely manipulate human behavior at the individual level (Bashivan et al., 2019; Mueller et al., 2023; Ponce et al., 2019; Shahbazi et al., 2024; Walker et al., 2019).

Here we examine whether individual differences in ANNs capture individual differences in human behavior. We trained and evaluated 60 instances from three ANN architectures on a digit recognition task and observed substantial variability in accuracy, confidence, and response time (RT). We then tested the alignment between each human subject and each ANN instance. We found that these individual differences in ANN instances mapped consistently onto the individual differences produced by 60 humans performing the same task, with the mapping strength often approaching the strength of the human-to-human benchmark. Remarkably, these human-ANN mappings were not confined to a single behavioral metric: ANN instances that mimic an individual in one behavioral metric also tended to mimic the same individual on other metrics. These results extended to a 10-choice blurry object recognition task, although the human-ANN mappings were less robust than the human-human benchmark in that task. Taken together, these findings open the possibility of using ANNs in uncovering the mechanisms underlying individual variability in human behavior.

## 2 Results

To explore whether the individual differences in ANNs are related to individual differences in humans, we first reanalyzed data from a previous study where N=60 human subjects performed an 8-choice noisy digit recognition task (Figure 1a; Rafiei et al., 2024). We recorded accuracy, confidence, and RT on each trial. We trained models based on three different ANN architectures – RTNet (Rafiei et al., 2024), AlexNet (Krizhevsky et al., 2012), and ResNet18 (He et al., 2015). For each architecture, we trained 60 independent instances by changing the random initialization. All models were trained to ceiling, achieving over 97% accuracy on a validation dataset of noiseless images. We then adjusted the noise level in the images separately for each network architecture, such that the average accuracy of the 60 instances from that architecture (RTNet: 0.71; AlexNet: 0.72; ResNet18: 0.71) matched the average accuracy of human subjects (0.71; Figure 1b). Note that this procedure simply adjusted the overall noise level of the images presented to the whole population of instances of a given architecture; we did not do anything to directly align individual human subjects to individual ANN instances.

**Figure 1.**
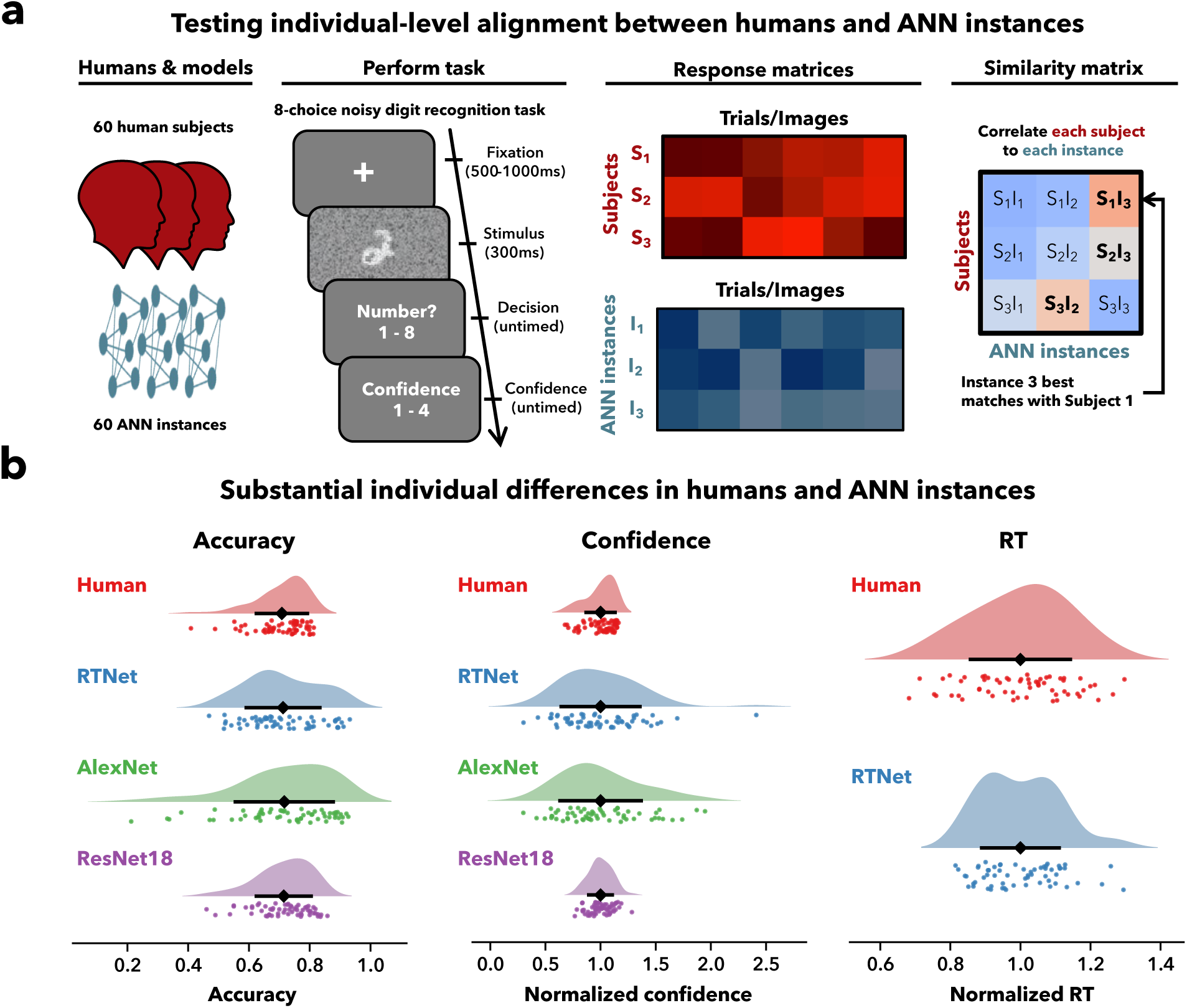
Task schematics and behavioral differences in humans and ANN instances. (a) Schematics of testing individual-level alignment. Human subjects (N=60) and ANN instances (N = 60 from each of three architectures: RTNet, AlexNet, ResNet18) performed an 8-choice noisy digit recognition task. On each trial, human subjects first saw a fixation cross for 500-1000 ms, followed by a noisy image of a handwritten digit from the MNIST dataset presented for 300 ms. Subjects then reported their decision, followed by their confidence rating. ANN instances were presented the same MNIST images and we extracted responses and confidence ratings equivalent to those in humans. Responses from human subjects and ANN instances were structured into response matrices. We correlated the image-by-image responses of each subject to each ANN instance to construct the similarity matrix, where each cell indicates how well each subject aligns to each ANN instance. (b) Substantial individual differences in humans and ANN instances. Confidence and RT are normalized by their mean across subjects or instances for visualization. Diamond markers denote the mean. Error bars show standard deviation. Dots represents individual subjects or individual ANN instances.

### 2.1 Substantial individual differences in humans and ANN instances

We first confirmed that there were substantial individual differences in both ANN instances and humans (Figure 1b). Indeed, even though ANN instances from each architecture differed only in their random initializations, they showed a very large range in average accuracy when tested on noisy images (RTNet: range = [0.47, 0.93], SD = 0.12; AlexNet: range = [0.21, 0.93], SD = 0.16; ResNet18: range = [0.46, 0.86], SD = 0.09). Surprisingly, the variability among ANN instances was even larger than the variability among human subjects (range = [0.41, 0.81], SD = 0.08). We obtained similar results for confidence and RT. Since confidence and RT are on different scales across humans and each ANN architecture, we quantified the individual variability as the coefficient of variation (CV, standard deviation divided by the mean). Using this metric, we observed greater individual variability in confidence for RTNet and AlexNet than in humans (humans: CV = 0.13; RTNet: CV = 0.36; AlexNet: CV = 0.37), whereas ResNet18 showed a similar level of variability to humans (ResNet18: CV = 0.11). RT was assessed only for RTNet (as the other models do not produce RT) and showed a comparable level of variability between humans and RTNet (humans: CV = 0.14; RTNet: CV = 0.11). Similar results were found when examining variability in the full trial-level response distributions using Wasserstein distances (Supplementary Figure 1 & Supplementary Results). These results demonstrate that ANNs that only differ in their random initialization exhibit large individual differences that either match or exceed the variability among human subjects.

### 2.2 Individual ANN instances vary in their alignment with individual human subjects

Having confirmed the existence of substantial individual differences in overall accuracy, confidence, and RT for ANNs and humans, we explored how well each ANN instance aligns with each human subject. For each of the three behavioral metrics, we computed the correlations between all pairs of human subjects and network instances. We found a wide range of correlations between ANN instances and individual humans (see Figure 2a for accuracy and Supplementary Figure 2 for confidence and RT). For example, when considering the correlations for accuracy only, the range of correlation values was [-0.02, 0.57] for human-human alignment, [-0.24, 0.48] for human-RTNet alignment, [-0.15, 0.48] for human- AlexNet, and [-0.01, 0.47] for human-ResNet18 correlations. Similar results were obtained for confidence (human-human: [-0.09, 0.55], human-RTNet: [-0.20, 0.47], human-AlexNet: [-0.22, 0.57]; human-ResNet18: [-0.08, 0.47]) and RT (human-human: [-0.04, 0.47], human-RTNet: [-0.21, 0.43]).

**Figure 2.**
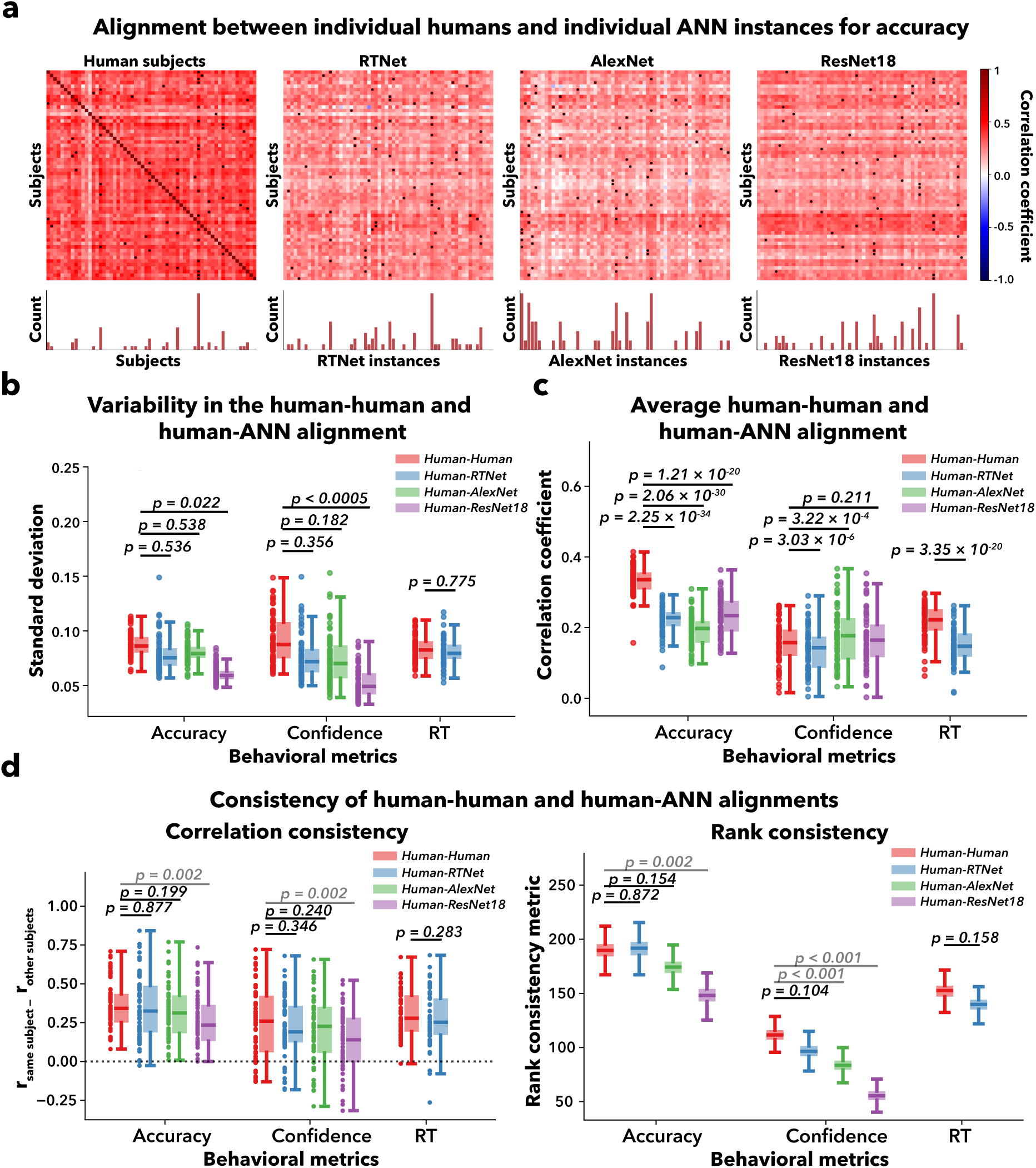
Individual ANN instances vary in their alignment with individual human subjects. (a) Similarity between each subject and each ANN instance in accuracy. We constructed the similarity matrix of 60 human subjects and the 60 instances of each ANN architecture, with higher values indicating a better alignment between an individual human and a specific ANN instance. The black dots indicate the model instance with the best fit for each subject (i.e., there is one black dot for each row). The histograms underneath show how often each subject or ANN instance was the best match across all human subjects. (b) Variability in the human-human and human-ANN alignment. The variability in alignment between individual humans and ANN instances was similar to the variability in alignment between each human subjects and the remaining subjects. P-values indicate the significance of pairwise differences in the standard deviation of the similarity matrix. (c) Average human-human and human-ANN alignment. Individual ANN instances showed slightly lower overall alignment with humans compared to the alignment between different humans. Dots show individual human subjects or ANN instances. P-values indicate the significance of pairwise differences in the average correlation. (d) Consistency of human-human and human-ANN mapping. Using both the correlation and rank methods of quantifying consistency converge on the same findings that individual ANN instances map to human subjects as consistently as humans do. This effect is true for RTNet and AlexNet, whereas ResNet18 shows lower consistency than the human benchmark. Dots show individual human subjects. Dotted horizontal line denotes zero consistency between the two maps. P-values for correlation consistency indicate the significance of pairwise differences in the group-adjusted correlation between human-human and human-ANN mapping, computed using paired t-tests. P-values for rank consistency indicate the significance of pairwise differences in the rank consistency metric. For all panels, boxes indicate the 25^th^ and 75^th^ percentiles, the central line denotes the median, and whiskers extend to 1.5 x IQR.

The similarity matrices in Figure 2a show that the same human subject or same ANN instance sometimes provides the best match to the data for multiple human subjects. We counted how often each subject or ANN instance was the best match across all human subjects and demonstrated largely similar patterns for the human-human and human-ANN alignment (Supplementary Figure 3 & Supplementary Results).

To further assess the amount of variation in the human-human and human-ANN alignment, we computed the standard deviation of the human-human and human-ANN correlation values for each of the 60 human subjects. This allowed us to examine, for each subject, the variability in how well other humans or ANN instances correlate with that subject. We found that variability in human-human correlations was broadly similar to the variability in human-ANN correlations (Figure 2b). Specifically, human-RTNet variability was indistinguishable from human-human variability for accuracy (*p* = 0.536), confidence (*p* = 0.356), and RT (*p* = 0.775). Human-AlexNet variability was also similar to human-human variability for both accuracy (*p* = 0.538) and confidence (*p* = 0.182). In contrast, human-ResNet18 correlations demonstrated significantly lower variability than human-human correlations for both accuracy (*p* = 0.022) and confidence (*p* < 0.0005), though the standard deviations decreased by a relatively small amount (up to 33%). Overall, the variability in how well individual humans are aligned with other individual humans was either comparable or only slightly higher than the variability in how well they are aligned with ANN instances.

For completeness, we also examined the strength of human-human and human-ANN alignment. We found that the average human-human correlations were higher than the average human-ANN correlations for accuracy (*p*’s < 0.001 for all three architectures; Figure 2c). This was also true for confidence and RT for RTNet (both *p’s* < 0.001). Notably, the average human-ANN correlations matched the human-human correlations for confidence in the case of ResNet18 (*p* = 0.211) and even exceeded the human-human correlations for AlexNet (*p* = 3.22 x 10^-4^). These results demonstrate that individual ANN instances are overall not as good as human subjects at predicting individual human performance, although the difference is relatively small (e.g., for RTNet the decrease in correlation was 33%, 13%, and 30% for accuracy, confidence, and RT, respectively).

### 2.3 Individual ANN instances show consistent mapping with human subjects across stimulus subsets

The results in the previous section demonstrated the existence of substantial individual differences between ANN instances in their alignment with individual human subjects. However, some amount of individual difference is expected from noise alone. Therefore, it is critical to demonstrate that the differences between ANN instances in how they align with individual subjects are consistent and persist even when assessed using separate sets of images. To this end, we performed 1000 iterations of randomly splitting the trials into two equal parts and computed the correlation matrix for each split. Finally, we evaluated the consistency of the correlation matrix and compared the obtained consistency values to human-human consistency as a benchmark.

We first quantified the consistency of the human-ANN similarity matrices by computing the Pearson correlation between the two halves of each of the 1000 trial splits within each individual subject. We then group-adjusted these values for each trial split by subtracting the average correlation between the same subject and all other subjects, which removes any correlations stemming from ANN instances that may perform uniformly well or uniformly badly in matching all human subjects. We found that human-RTNet correlations were as consistent as the human-human benchmark for accuracy (*t*(59) = 0.155, *p* = 0.877, BF_01_ = 5.08; Figure 2d), confidence (*t*(59) = 0.946, *p* = 0.346, BF_01_ = 3.44), and RT (*t*(59) = 1.08, *p* = 0.283, BF_01_ = 3.04). The same was also true for AlexNet (accuracy: *t*(59) = 1.29, *p* = 0.199, BF_01_ = 2.42; confidence: *t*(59) = 1.18, *p* = 0.240, BF_01_ = 2.74) but not for ResNet18, which exhibited lower consistency than the human-human benchmark for both accuracy (*t*(59) = 3.10, *p* = 0.002, BF_10_ = 13.3) and confidence (*t*(59) = 3.17, *p* = 0.002, BF_10_ = 16.4).

One concern with using Pearson correlation is that high consistency values could be driven by a few ANN instances that always show bad alignment with most subjects and it could be that our subtraction method doesn’t fully correct for this effect. To mitigate this concern, we conducted a complementary analysis. For each subject, we computed the rank order of how well each ANN instance (or each subject except the selected one) is aligned with that subject. Then, for each pair of subjects, we calculated the difference in the ranks of ANN instances (or other subjects) for each of two subsets of the data. We then took the average of the product of these differences across the two subsets, with positive values indicating consistent rank ordering. Using this alternative metric of consistency, we found that human- RTNet similarity matrix had comparable consistency to the human-human benchmark for accuracy (*p* = 0.872, 95% bootstrap CI = [-24.6, 21.6]), confidence (*p* = 0.104, 95% bootstrap CI = [-2.92, 32.0]), and RT (*p* = 0.158, 95% bootstrap CI = [-5.03, 31.1]). Similarly, human-AlexNet similarity matrix was as consistent as the human benchmark for accuracy (*p* = 0.154, 95% bootstrap CI = [-5.10, 35.6]), but not for confidence (*p* < 0.001, 95% bootstrap CI = [11.6, 44.5]). Lastly, the human-ResNet18 similarity matrix exhibited significantly lower consistency than the human benchmark for both accuracy and confidence (both p’s < 0.003). Overall, both the Pearson correlation and the rank order methods of quantifying consistency demonstrate that individual ANN instances for RTNet and AlexNet map to human subjects as consistently as human subjects map among themselves (while ResNet18 instances show lower consistency). These results demonstrate that the variability in human-ANN mapping is not simply due to noise but reflects meaningful variability among ANN instances.

### 2.4 ANN instances that mimic an individual on one metric also mimic the same individual on other metrics

Our results so far show that a given ANN instance is more similar to some human subjects than others. However, we showed this fact separately for each of our three behavioral metrics (accuracy, confidence, and RT). Here we test whether an ANN instance that mimics a subject’s behavior on one metric (e.g., accuracy) also captures that subject’s behavior for other metrics (e.g., confidence). To this end, we computed the correlations between the similarity matrices for each subject across pairs of behavioral metrics (Figure 3a). Importantly, this analysis assesses the mapping between the two similarity matrices built using pairs of behavioral metrics rather than directly correlating the behavioral metrics themselves. We found that the human-ANN across-metric mapping were significantly positive for all pairs of behavioral metrics, including accuracy-confidence (human-RTNet: average *r* = 0.340; *t*(59) = 11.8, *p* = 4.25 x 10^-17^; human-AlexNet: average *r* = 0.367; *t*(59) = 15.7, *p* = 1.14 x 10^-22^; human-ResNet18: average *r* = 0.109; *t*(59) = 4.14, *p* = 1.14 x 10^-4^; Figure 3b), accuracy-RT (human-RTNet: average *r* = 0.333; *t*(59) = 11.6, *p* = 6.78 x 10^-17^), and confidence-RT (human-RTNet: average *r* = 0.440; *t*(59) = 12.9, *p* = 8.48 x 10^-19^). Moreover, the human-RTNet across-metric mappings were statistically indistinguishable from the human-human control for accuracy-confidence (*t*(59) = −1.68, *p* = 0.098, BF_01_ = 1.88) and accuracy-RT (*t*(59) = −0.71, *p* = 0.480, BF_01_ = 5.56), and significantly outperformed the human-human benchmark for RT-confidence (*t*(59) = 5.23, *p* = 2.33 x 10^-6^, BF_10_ = 7.14 x 10^3^). Similarly, the human-AlexNet mapping was comparable to the human-human benchmark for accuracy-confidence (*t*(59) = −1.02, *p* = 0.313, BF_01_ = 4.33). In contrast, the human-ResNet18 mapping was significantly worse than the human-human benchmark for accuracy-confidence (*t*(59) = −9.36, *p* = 2.87 x 10^-13^, BF_10_ = 2.76 x 10^10^).

**Figure 3.**
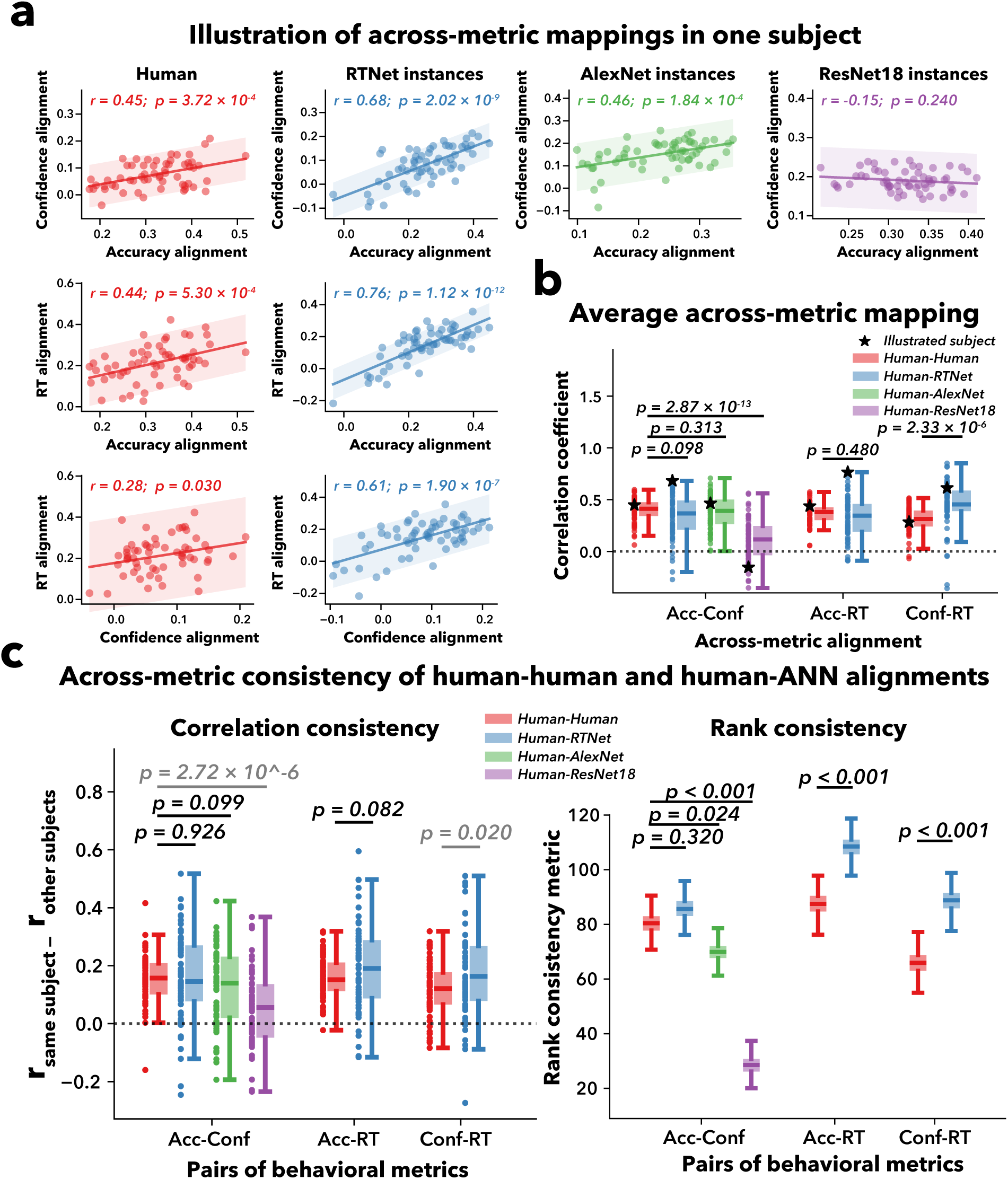
ANN instances that mimic an individual on one metric also mimic the same individual on other metrics. (a) Scatterplots for one example human subject showing how alignment with another human or ANN instance on one metric relates to alignment with the same human or ANN instance on another metric. Dots show individual ANN instances (or all other subjects). (b) Average strength of the across-metric mappings. The human-ANNs across-metric mapping showed comparable or even higher correlation than the human-human benchmark, except for human-ResNet18, which showed lower correlation. Dots show individual human subjects. Dotted horizontal line denotes a complete lack of across-metric mapping. P-values indicate the significance of pairwise differences in average across-metric mapping of all subjects. (c) Across-metric consistency of human-human and human-ANN alignments. We assessed across-metric consistency using the same methods previously applied to quantify within-metric consistency. Both methods of quantifying consistency yielded similar results: individual ANN instances map to human subjects as consistently as humans do, which is true for both RTNet and AlexNet, whereas ResNet18 shows lower consistency than the human-human benchmark. Dots show individual human subjects. Dotted horizontal line denotes zero across-metric consistency between two maps. P-values for correlation consistency indicate the significance of pairwise differences in the group-adjusted correlation between human-human and human-ANN alignment, computed using paired t-tests. P-values for rank consistency indicate the significance of pairwise differences in the rank consistency metric. For all panels, boxes indicate the 25^th^ and 75^th^ percentiles, the central line denotes the median, and whiskers extend to 1.5 x IQR.

As before, we confirmed that these across-metric mappings reflect robust relationships by evaluating their consistency over 1000 iterations of random splits of the data (Figure 3c). Critically, across-metric consistency was assessed by comparing split 1 of one metric to split 2 of another, which ensures that the observed consistency cannot be attributed to shared data variance. Using Pearson correlation, we found that the human-RTNet across-metric consistency was either similar to or higher than the human-human benchmark in all pairs of behavioral metrics (accuracy-confidence: *t*(59) = −0.093, *p* = 0.926, BF_01_ = 5.13; accuracy-RT: *t*(59) = −1.76, *p* = 0.082, BF_01_ = 1.28; RT-confidence: *t*(59) = −2.37, *p* = 0.020, BF_10_ = 2.36). The same was true for the human-AlexNet accuracy-confidence consistency (*t*(59) = 1.66, *p* = 0.099, BF_01_ = 1.48), whereas human-ResNet18 showed significantly lower consistency than to the human-human benchmark (*t*(59) = 4.97, *p* = 2.72 x 10^-6^, BF_10_ = 6.68 x 10^4^). Similar results were also obtained using rank order. However, using the rank metric, human-RTNet exhibited significantly greater consistency compared to the human-human benchmark in the accuracy-RT (*p* < 0.001, 95% bootstrap CI = [-32.1, − 10.9]) and RT-confidence relationships (*p* < 0.001, 95% bootstrap CI = [-33.8, −12.0]), and matched the human-human benchmark in accuracy-confidence consistency (*p* = 0.320, 95% bootstrap CI = [-15.7, 4.55]). Human-AlexNet and human-ResNet18, however, demonstrated significantly lower consistency than to the human-human benchmark in the accuracy-confidence pair (AlexNet: *p* < 0.001, 95% bootstrap CI = [1.39, 21.0]; ResNet18: *p* < 0.001, 95% bootstrap CI = [42.1, 62.4]). These results show that the correspondence between individual differences in humans and ANN instances extends even across metrics, suggesting that an ANN instance may mimic an individual on a deeper level than what is reflected by a single metric.

### 2.5 Human-ANN mapping is not simply due to performance variability in ANNs and humans

Thus far, we demonstrated that ANN instances differing in their random initialization exhibit substantial individual differences that correspond to human individual differences across data subsets and behavioral metrics. However, it remains possible that the observed correspondence between ANN instances and human subjects is mostly driven by the variability in performance among ANN instances and human subjects. Specifically, both ANN instances and humans differed in their overall accuracy on the task, giving rise to individual differences in behavioral performance. Thus, ANN instances with higher or lower accuracy could preferentially map onto human subjects with correspondingly higher or lower performance. Such correspondence would suggest that consistent human-ANN mappings may simply reflect similarities in overall performance rather than genuine instance-specific behavioral similarities. Here we addressed this question in two different ways.

#### 2.5.1 Individual ANN instances show consistent mapping with human subjects across different levels of average performance

We first examined how the mapping between humans and ANNs varied across neural network populations spanning different levels of performance. Particularly, in our current evaluation, we adjusted the noise level of the stimulus to calibrate the overall performance of the ANN population to that of human subjects. Here, for each of the three architectures (RTNet, AlexNet, and ResNet18), we used five levels of stimulus noise so that average accuracy of the ANN population was equal to *mean* − 2*SD*, *mean* − 1*SD*, *mean*, *mean* + 1*SD*, and *mean* + 2*SD* of the human performance distribution (the exact targets were 51.8%, 61.6 %, 71.4 %, 81.2%, and 91.0%). We then repeated the same mapping analyses for each stimulus noise level.

We found that in all three architectures, all stimulus noise levels led to robustly positive mapping consistency (all *p*’s < 0.001, Figure 4a), suggesting that the presence of mapping between humans and ANN instances despite differences in average performance. We next examined how the human-ANN mapping consistency varied across the five stimulus noise levels. We found that the strengths of the human-ANN mappings were largely similar across the three highest-noise stimuli (Figure 4a). However, the lowest two stimulus noise levels led to lower mapping consistency for accuracy that was significantly below the human-human benchmark (all *p*’s < 0.01). This was likely due to a ceiling effect: as population-level performance increases, behavioral performance becomes increasingly compressed toward the upper bound, reducing the magnitude of individual differences and thus lowering the consistency of the human-ANN mapping. Note that this pattern was generally absent in confidence where no similar ceiling exists, though it also occurred for RT in RTNet. This general pattern was also observed for across-metric mapping consistency (Supplementary Figure 4). These findings indicate that the human-ANN mappings are generally consistent across different levels of overall ANN performance and do not only appear when performance is matched across humans and ANNs.

**Figure 4.**
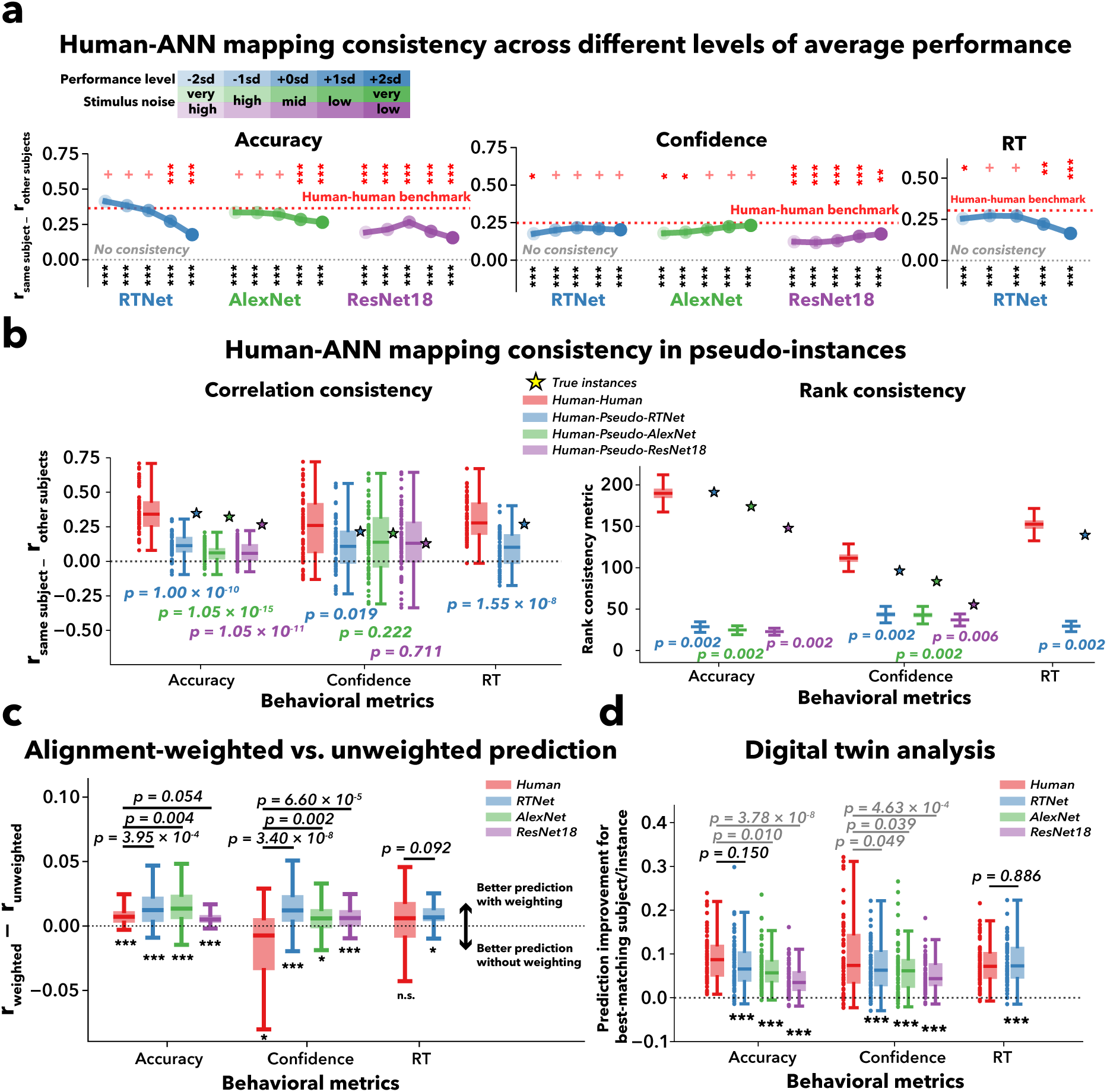
Control analyses and predicting individual human behavior. (a) Human-ANN mapping consistency across five levels of average performance. We used five levels of stimulus noise so that average accuracy of the ANN population was equal to ***mean*** − **2*SD***, ***mean*** − **1*SD***, ***mean***, ***mean*** + **1*SD***, and ***mean*** + **2*SD*** of the human performance distribution. The results show that the individual human-ANN mappings are generally consistent across different levels of overall ANN performance. Grey and red dotted lines denote the absence of mapping consistency and the human-human mapping benchmark, respectively. Black asterisks indicate significance against zero. Red asterisks indicate pairwise-comparisons with the human-human benchmark. (b) Human-ANN mapping consistency in pseudo- instances. We created “pseudo-instances” from a single ANN instance by systematically varying stimulus noise. The lower mapping consistency observed for pseudo-instances than true instances demonstrates that performance similarity alone cannot fully account for the correspondence we observed between individual human subjects and true ANN instances. P-values indicate the significance of pairwise differences between the true instances (Star) and pseudo-instances. (c) Alignment-weighted improvement among humans and three ANN architecture. We used the correlation in the similarity matrix as weights to predict human behavior in a held-out data subset. This alignment-weighted prediction was then compared to an unweighted average of all ANN instances. All ANNs showed significant improvement in prediction with the alignment weighting and even outperformed the human benchmark. (d) Digitial twin prediction analysis. We identify the best-matching neural network instances (digital twin) and predict human behavior in a held-out data subset. All ANNs showed significant improvement in prediction with the digital twin than the predictions from all other instances. Dot shows individual subjects. For all panels, unless stated otherwise, boxes indicate the 25^th^ and 75^th^ percentiles, the central line denotes the median, and whiskers extend to 1.5 x IQR. P-values above the bars indicate the significance of pairwise differences between humans and ANNs in prediction improvement. Black asterisks below the dotted line indicate significance for one-sample t-tests against zero. *, *p* < 0.05; **, *p* < 0.01; ***, *p* < 0.001; n.s. or + not significant, *p* > 0.05.

#### 2.5.2 Pseudo-instances exhibit substantially lower human-ANN mapping strengths

The results in the previous section suggested that the mapping consistency between individual subjects and ANN instances is broadly robust across different levels of average performance. To further demonstrate that human-ANN mappings do not simply reflect similarities in overall performance, we have to dissociate instance-specific differences in ANNs from differences in overall performance. To this end, we created “pseudo-instances” from a single ANN instance by systematically varying the level of stimulus noise during testing. Specifically, for each of the 60 ANN instances in one architecture, we took an ANN instance and generated 60 pseudo-instances by testing this same instance under different levels of noise, thereby producing a range of performance levels across pseudo-instances while keeping the underlying ANN instance identical. We then repeated the same mapping analyses for each set of 60 pseudo-instances, resulting in 60 independent sets of analyses corresponding to the 60 original ANN instances. If the individual human-ANN map is simply driven by overall performance similarities, we would expect these pseudo-instances to show the same mapping consistency as the true ANN instances.

We found that the true instances showed significantly higher mapping consistency than their pseudo- instances counterparts in RTNet for all three behavioral metrics and for both the correlation and rank analyses (all *p*’s < 0.02, Figure 4b). AlexNet and ResNet18 also showed greater mapping consistency in true instances than pseudo-instances for accuracy in both the correlation and rank analyses (all *p*’s < 0.001) but for confidence this effect was only significant for the rank analyses (p’s < .006 for the two rank analyses; *p*’s > 0.2 for the two correlation analyses). These results indicate that the mapping between individual differences in humans and ANN instances is driven by instance-specific factors beyond similarities in overall performance. However, it is also important to note that the mapping consistencies for all pseudo-instances were significantly larger than zero across all behavioral metrics and ANN architectures (all *p*’s < 0.001). We found similar results for across-metric mapping consistency (Supplementary Figure 5). These findings indicate that similarities in overall performance do contribute to the human-ANN mapping. However, the lower mapping consistency observed for pseudo-instances demonstrates that performance similarity alone cannot fully account for the strong correspondence we observed between individual human subjects and true ANN instances.

### 2.6 Leveraging individual differences in ANN instances to predict single-subject responses

We next explored whether we can leverage the mappings between ANN instances and human subjects to improve the prediction of individual human behavior. Specifically, if the individual differences in ANN instances truly reflect the individual differences in humans, the human-ANN mapping, which captures human individual differences, should improve prediction of individual human behavior in held-out data compared with approaches that do not account for individual differences. To test this hypothesis, we computed the human-ANN similarity matrix from one data subset to learn the alignment strength between each subject and ANN instance. We then used the correlations in the similarity matrix as weights to predict behavior in a held-out data subset. This procedure weighs the ANN instances in proportion to how similar they are to a given human subject. Finally, we compared this alignment- weighted prediction with an unweighted prediction that simply took averages of all ANN instances.

We evaluated the prediction performance within each behavioral metric (Figure 4c). All three ANNs showed significantly better alignment-weighted prediction compared to the unweighted prediction. This was true for accuracy (RTNet: *t*(59) = 5.47, *p* = 9.67 × 10^-7^; AlexNet: *t*(59) = 4.98, *p* = 5.86 × 10^-6^; ResNet18: *t*(59) = 5.89, *p* = 2.00 × 10^-7^), and confidence (RTNet: *t*(59) = 5.33, *p* = 1.60 × 10^-6^; AlexNet: *t*(59) = 2.06, *p* = 0.044; ResNet18: *t*(59) = 5.39, *p* = 1.27 × 10^-6^), and RT (RTNet: *t*(59) = 2.50, *p* = 0.015). While these improvements were modest (maximum improvement of 0.017 in correlation), they were remarkably consistent across different subjects. As a benchmark, we used the same procedure to assess the difference between human alignment-weighted prediction and unweighted prediction in held-out data for each subject (note that this comparison doesn’t consider the raw prediction accuracy for each type of prediction, which was higher for humans than ANNs; Supplementary Figure 6). We found that both RTNet and AlexNet outperformed the human benchmark for accuracy (RTNet: *t*(59) = 3.76, *p* = 3.95 × 10^-4^; AlexNet: *t*(59) = 2.99, *p* = 0.004) and confidence (RTNet: *t*(59) = 6.35, *p* = 3.48 × 10^-8^; AlexNet: *t*(59) = 3.18, *p* = 0.002), but RTNet matched the human benchmark for RT (RTNet: *t*(59) = 1.71, *p* = 0.092). In contrast, ResNet18 showed a larger improvement than in the human benchmark for confidence (*t*(59) = 4.30, *p* = 6.60 × 10^-5^) but not for accuracy (*t*(59) = 1.97, *p* = 0.054).

The above approach validated that incorporating alignment weights in prediction could improve prediction of single-subject responses. Yet, this approach predicts single-subject responses with an ensemble of network instances, with each instance contributing to the prediction according to its similarity to the subject. An alternative approach of predicting single-subject responses is the *digital twin* approach (Wood et al., 2024), which identifies the best-matching instance for one subject, and uses that instance as a digital twin for prediction. Here we employed this approach by first identifying the best-matching ANN instance for each subject using half of the data. We then predicted each subject’s responses in the other half of the data using only the responses from their best-matching ANN instance (i.e., their “digital twin”). We then compared these predictions with those obtained using instances other than the best-matching instance for each subject separately for accuracy, confidence, and RT.

We found that for all three ANN architectures, the digital twin predicted a subject’s behavior better than the average of all other instances (all *p*’s < 0.001; Figure 4d; see Supplementary Figure 7 for the raw prediction accuracy). Critically, we also compared these results with the human-human benchmark. Specifically, for each subject, we identified the best-matching other subject and used their responses to predict the target subject’s responses. We then compared these predictions with those obtained using all other subjects. We found that the improvement in prediction for the digital RTNet twin was largely comparable to the improvement for the best-matching human across the three behavioral metrics (Accuracy: *t*(59) = 1.45, *p* = 0.150; BF_01_ = 2.0; Confidence: *t*(59) = 2.00, *p* = 0.049; BF_01_ = 0.86; RT: *t*(59) = − 0.144, *p* = 0.886 ; BF_01_ = 5.10). However, the digital-twin improvements for both AlexNet and ResNet18 were significantly smaller than the corresponding best-matching-human benchmarks for both accuracy and confidence (all *p*’s < 0.04). Together, these results demonstrate the ability of leveraging individual differences in ANNs to predict individual human behavior.

### 2.7 Extension to 10-choice object recognition

The results above were obtained using a relatively simple stimulus space – hand-written digits. Therefore, we further tested how well these results extend to a more complex stimulus space comprised of naturalistic objects. Generalizing to a complex object space comes with several caveats. First, unlike for hand-written digits, ANN models are harder to train to near-perfect accuracy for common object datasets. Thus, matching average human and ANN accuracy requires substantially lower image degradation for ANNs compared to humans. Second, the added complexity of object space (compared to digits space) means that we are likely to observe lower average alignments between humans and ANNs. Given these factors, ANN instances are unlikely to fully capture the variability observed in human behavior when testing on objects. Nonetheless, we examined whether individual differences across ANN instances could still account for some components of human individual variability.

To this end, we conducted a preregistered experiment where N=60 human subjects performed a 10-choice blurry object recognition task (Figure 5a) and recorded accuracy, confidence, and RT on each trial. As with our MNIST analyses, we trained 60 unique instances of RTNet (Rafiei et al., 2024), AlexNet (Krizhevsky et al., 2012), and ResNet18 (He et al., 2015) by changing the random initialization. All model instances were trained in identical fashion and achieved at least 75% accuracy on a validation dataset of images with no blurring. We then adjusted the blur level in the images separately for each network architecture, so that at the population-level, the average accuracy of the 60 ANN instances from each architecture matched the average accuracy of the 60 human subjects.

**Figure 5.**
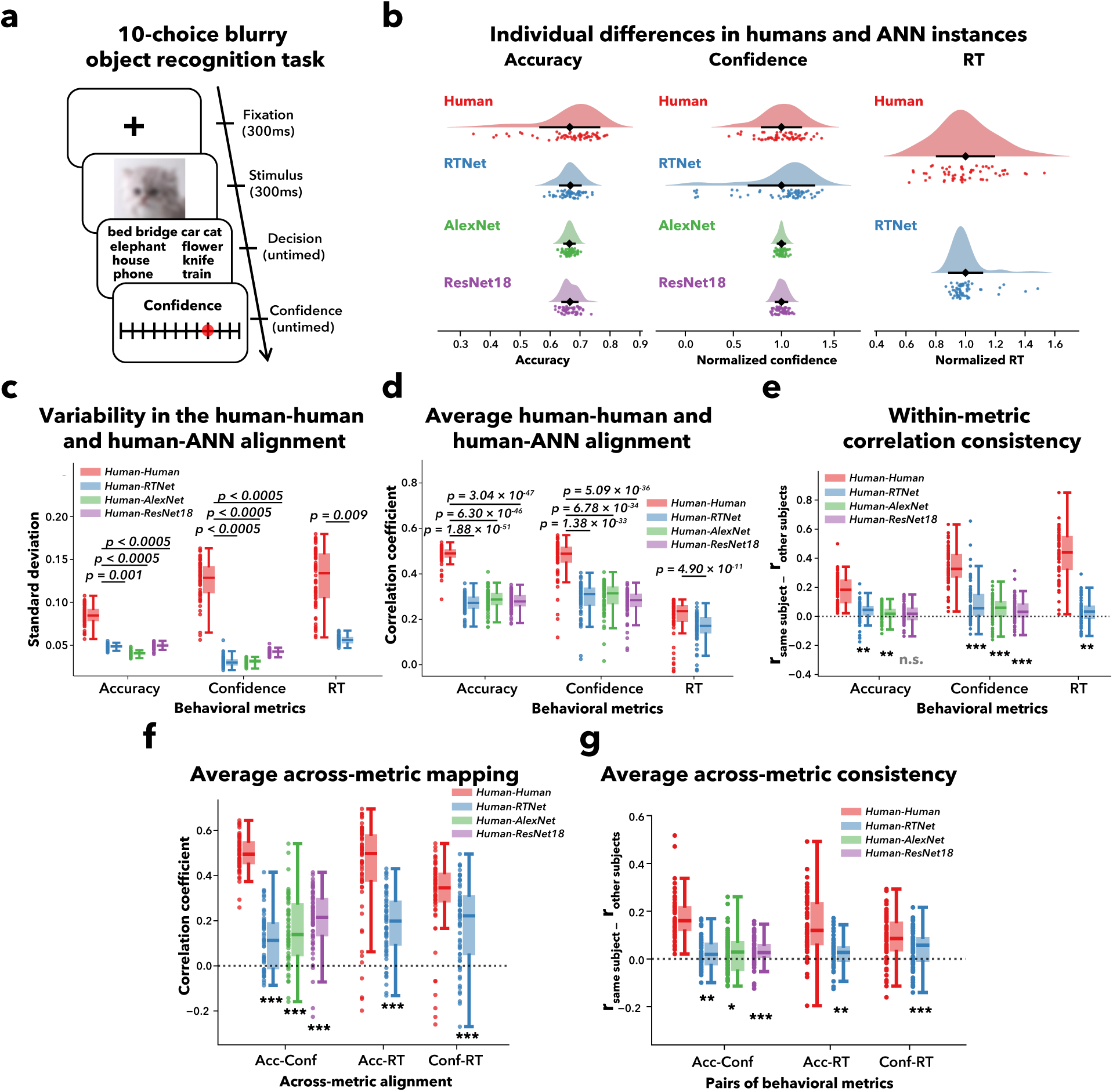
Extension to the 10-choice object recognition task. (a) Task schematics for the 10-choice blurry object recognition task. On each trial, subjects saw a blurred image of an object from the EcoSet dataset and reported their decision, followed by their confidence rating. (b) Individual differences in humans and ANN instances. Confidence is normalized by the mean across subjects or instances for visualization. Dots represent individual subjects or ANN instances. Unlike the 8-choice task, we found that humans demonstrate significantly more individual variability than ANN instances. (c) Variability in the human-human and human-ANN alignment. The variability in how well individual humans are aligned with other individual humans is significantly higher than the variability in how well they are aligned with ANN instances. P-values indicate the significance of pairwise differences in the standard deviation of the similarity matrix, computed using bootstrap resampling with 2000 data subsets. (d) Average human-human and human-ANN alignment. Individual ANN instances showed significantly lower overall alignment with humans compared to the alignment between different humans. P-values indicate the significance of pairwise differences in the average correlation. (e) Within-metric correlation consistency of human-human and human-ANN alignments. Most human-ANN alignments demonstrate significantly higher consistency than what is expected from chance, although lower than the human-human benchmark. Asterisks below the bars indicate significance for one-sample t-tests against zero. (f) Average across-metric mapping. All human-ANNs across-metric correlations are significantly higher than zero. (g) Across-metric correlation consistency of human-human and human-ANN mappings. All human-ANN mappings demonstrate significantly better across-metric consistency than chance, although lower than the human-human benchmark. For all panels, unless otherwise noted, dots show individual human subjects, dotted lines denote zero consistency between two maps. Boxes indicate the 25^th^ and 75^th^ percentiles, the central line denotes the median, and whiskers extend to 1.5 x IQR. Asterisks below the bars indicate significance for one-sample t-tests against zero (*, *p* < 0.05; **, *p* < 0.01; ***, *p* < 0.001; n.s., not significant).

We first examined whether there were substantial individual differences in both ANN instances and humans (Figure 5b). Unlike the 8-choice digit task, where ANN instances and humans showed comparable behavioral variability, ANN instances exhibited markedly reduced variability across instances compared to human subjects for both accuracy (human SD = 0.098, RTNet SD = 0.034, AlexNet SD = 0.017, ResNet18 SD = 0.025) and RT (human CV = 0.191, RTNet CV = 0.111). An exception was confidence, for which RTNet (CV = 0.340) exhibited higher variability than humans (CV = 0.204), while AlexNet (CV = 0.038) and ResNet18 (CV = 0.058) showed substantially reduced variability. Similar results were obtained using Wasserstein distances (Supplementary Figure 8 & Supplementary Results).

We next computed the variations in the human-ANN and human-human similarity matrices. Again, diverging from the 8-choice digit task results, variability in the human-ANN alignments was significantly lower than in the human-human counterparts for both accuracy and confidence (all *p’s* < 0.0005; Figure 5c). We also found that the average strength of human-human correlations was significantly higher than human-ANN correlations across all three network architectures and behavioral metrics (all *p’s* < 0.001; Figure 5d). These results demonstrate that in the 10-choice object recognition task, individual ANN instances that differ in random initialization produce less variability and weaker alignment than human subjects themselves in predicting individual human performance.

To assess the robustness of the human-ANN mappings, we examined the consistency between human-human and human-ANN alignments across two subsets of the data. We found that human-RTNet alignment showed above-chance consistency for all three behavioral metrics (accuracy: *t*(59) = 3.29, *p* = 0.002; confidence: *t*(59) = 5.09, *p* = 3.96 × 10^-6^; RT: *t*(59) = 3.07, *p* = 0.003; Figure 5e). The same was true for human-AlexNet alignment (Accuracy: *t*(59) = 3.17, *p* = 0.002; Confidence: *t*(59) = 4.28, *p* = 7.07 × 10^-5^), but not for human-ResNet18 alignment, which demonstrated above-chance consistency only for confidence (*t*(59) = 4.19, *p* = 9.58 × 10^-5^) but not for accuracy (*t*(59) = 1.92, *p* = 0.060). Even though human-ANN alignments were generally consistent, that consistency was significantly lower than the human-human benchmark for all three behavioral metrics (all *p’s* < 0.001). Similar results were obtained using rank order consistency (Supplementary Figure 9a).

We then assessed the human-human and human-ANN alignments across pairs of behavioral metrics (accuracy, confidence, and RT). We found the human-ANN across-metric correlations were significantly positive for all pairs of behavioral metrics, including accuracy-confidence (human-RTNet: average *r* = 0.110; *t*(59) = 6.80, *p* = 5.83 x 10^-9^; human-AlexNet: average *r* = 0.155; *t*(59) = 7.79, *p* = 1.23 × 10^-10^; human- ResNet18: average *r* = 0.200; *t*(59) = 11.3, *p* = 1.87 × 10^-16^; Figure 5f), accuracy-RT (human-RTNet: average *r* = 0.184; *t*(59) = 9.92, *p* = 3.47 × 10^-14^), and confidence-RT (human-RTNet: average *r* = 0.185; *t*(59) = 7.68, *p* = 1.94 × 10^-10^). However, all these human-ANN across-metric correlations were significantly lower than the human-human benchmark (all *p’s* < 0.001). We further examined the consistency of these across-metric mappings. We found that the human-ANN across-metric correlation showed significantly higher consistency than chance in all pairs of behavioral metrics and for all network architectures, including RTNet (accuracy-confidence: *t*(59) = 3.15, *p* = 0.003; accuracy-RT: *t*(59) = 2.90, *p* = 0.005; RT-confidence: *t*(59) = 4.16, *p* = 1.05 × 10^-4^; Figure 5g), AlexNet (accuracy-confidence: *t*(59) = 2.14, *p* = 0.036) and ResNet18 (accuracy-confidence: *t*(59) = 4.16, *p* = 1.06 × 10^-4^). Yet, the strengths of these mappings were lower than the human-human benchmark for all three pairs of behavioral metrics (all *p’s* < 0.001). Similar results were also obtained using rank order (Supplementary Figure 9b). As in the earlier analysis, we also tested whether leveraging individual differences across ANN instances could improve the prediction of individual human responses using both the alignment-weighting (Supplementary Figure 10) and digital twin approaches (Supplementary Figure 11). Both approaches showed improved predictive performance, although performance remained below the human benchmark. Together, these results show that individual ANN instances failed to fully mimic the variability observed in human subjects, but they do reliably capture some patterns across all three behavioral metrics, revealing a meaningful, albeit partial, mimicking of human behavioral variability.

## 3 Discussion

We investigated whether the variability across ANN instances trained with different weight initialization captures the individual differences observed in human behavior. Across two experiments, we demonstrated substantial behavioral variability between ANN instances. Critically, individual differences in ANN instances reliably captured the patterns of human behavioral variability. The mapping between ANN instances and individual humans generalized not only across different image subsets but also across multiple behavioral metrics (accuracy, confidence, and RT). Notably, the mapping was particularly strong for the digit recognition task, approaching the strength of human-to-human mapping benchmarks. Overall, these results reveal that the variability in ANN instances can serve as a proxy for human behavioral variability.

One of the central goals of NeuroAI is to build neural networks that behave like humans. Efforts in this space have shown, for instance, that aligning the visual diet of neural networks with the developmental trajectory of the human visual system produces more human-like behavior, including reduced texture bias and improved robustness to visual noise (Jang & Tong, 2024; Jinsi et al., 2023; Lu et al., 2025; Vogelsang et al., 2024). Most studies in this area, however, have focused on alignment at the population level, effectively tuning ANNs to resemble the behavior of an *average* human observer. In parallel, several recent papers have demonstrated that seemingly minor factors such as weight initialization (Mehrer et al., 2020), the order of the training images (Chow & Palmeri, 2024), and data augmentation (Jordan, 2024) can drive substantial variability in ANN instances within the same architecture. This work, however, primarily explored variability in internal representations, leaving open how such instance-to-instance differences relate to individual differences in human behavior. Our work extends and bridges these two lines of work by demonstrating substantial individual differences in neural networks across three behavioral metrics (accuracy, confidence, RT) and by directly evaluating human-ANN alignment at the individual level, establishing that individual differences in neural networks are not purely stochastic but can capture meaningful aspects of human behavioral variability.

Our work introduces a largely overlooked dimension of building human-like ANN systems – the ability to model individual differences among humans. Specifically, we propose that a truly human-like neural network should not only exhibit strong population-level alignment with the *average* human behavior, but also capture the structured idiosyncrasies that characterize individual humans. One way to conceptualize this distinction between average alignment and structured idiosyncrasies is in terms of the *solution space*, where the center corresponds to the average behavior, and the shape reflects structured idiosyncrasies (Figure 6). From this perspective, existing work focuses primarily on bringing the center of the ANN solution space closer to the center of the human solution space. This approach, however, overlooks the fact that the human solution space has a specific structure that may or may not match the structure of the ANN solution space. As progress in building human-like neural networks has begun to plateau (Conwell et al., 2024; Feather et al., 2025; Linsley et al., 2025; McNeal et al., 2024; Ratan Murty et al., 2021), we need new criteria for human-like neural networks that go beyond average alignment. Importantly, our data suggest that the structural alignment of individual differences is dissociable from average alignment. Specifically, although the three architectures we tested (RTNet, AlexNet, ResNet18) exhibited similar average alignment to humans (Figure 2c), there was a clear hierarchy of how well the ANNs captured the structured idiosyncrasies, with RTNet performing the best and ResNet18 performing the worst. These results highlight the need to explicitly evaluate and target the structure of individual differences when building human-like models.

**Figure 6.**
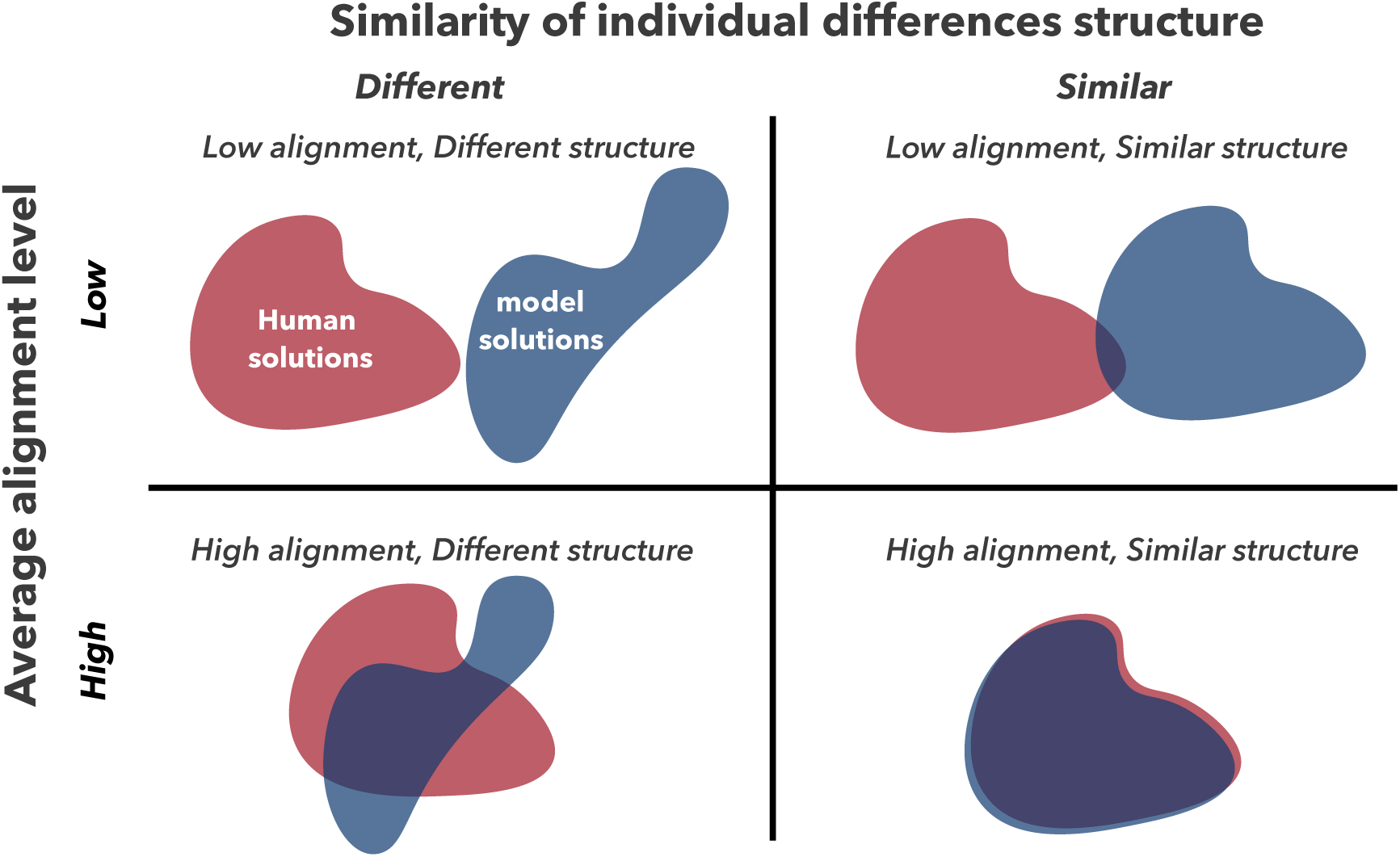
Illustration of the distinction between average alignment and structured individual differences. The 2 x 2 matrix illustrates the four possible scenarios in evaluating whether ANNs are human-like. Most existing work in NeuroAI focuses on average alignment by seeking to bring human solutions closer to the neural network solutions. We propose that a truly human-like neural network should not only exhibit strong average alignment with humans but also capture the structure of individual differences that characterize individual humans.

Although achieving high average alignment and a similar structure of individual differences are dissociable among different ANN architectures trained on the same task, the two objectives may be linked at the task level. Specifically, ANNs may be less human-like on some tasks than others, which would manifest in average alignment and quality of individual differences mapping changing in tandem across tasks. In our data, we observed both excellent average alignment and excellent individual differences mapping between humans and ANNs in the digit recognition task, whereas both these measures of how human-like a network is were reduced in the object recognition task. Future studies should explore a larger space of tasks to further probe whether average alignment and individual differences mapping vary together across tasks.

An open question that arises from our current study is why the ANN models were less human-like in the object recognition task. We speculate that the reason has to do with our ability to train the models to near-perfect level for undistorted images. Unlike the digit recognition task where ANNs can be easily trained to achieve near-ceiling performance for noiseless images, the object recognition task is too complex (and we lack a sufficient number of training images) to achieve such near-perfect accuracy for images with no blur (typical range = 75–85% accuracy). One consequence is that because humans are near perfect for undistorted images in the object recognition task, matching accuracy between humans and ANNs necessitates the use of lower image distortion for the ANNs compared to humans. Perhaps due to this difference in average image distortion, ANNs also exhibited substantially lower variability than humans in the object recognition task for all metrics (accuracy, confidence, RT). If this explanation is correct, it would suggest that human-like individual differences for a given task would emerge more strongly when ANNs can perform at or above the human level for undistorted images in that task.

Our results also provide a potential new approach to study the origins of human individual differences in visual perception. Rather than attempting to directly specify the mechanism that produced a given behavioral pattern in a particular individual, an alternative approach is to use multiple processes to generate populations of ANN instances. For instance, one can do so by manipulating random weight initialization (Mehrer et al., 2020), the order of the training images, the frequencies of items and categories during training (Chow & Palmeri, 2024), and data augmentation (Jordan, 2024). We can then ask which of these processes generated ANN instances that most closely resemble the variability observed in humans, with the best-matching process becoming a leading hypothesis for the origin of human variability in the tested task. Thus, ANNs may offer a tractable model system to study the origin of human visual variability.

This perspective of learning the computational processes that produce human-like behavioral distributions also raises the question of why random initialization produces individual differences in ANNs that map well onto individual differences in humans in our study. One possibility is that random weight initialization in neural networks loosely resembles the early sources of variability in human brain development. Critically, this mapping does not emerge in untrained networks (Supplementary Figure 12), suggesting that random initialization alone is insufficient to produce human-like individual differences and that training is critical for shaping these differences into a behaviorally meaningful solution space. In humans, these early sources of variability can include anatomical variability in neuronal organization and connectome structure that emerge prenatally (Gao et al., 2014; Xu et al., 2019) and functional stochasticity that may be generated by retinal waves and spontaneous cortical activity prior to any visual experience (Ackman et al., 2012; Hanganu et al., 2006; Wong, 1999). These factors introduce early variability in the developing human visual system, which could contribute to shaping the human solution space as individuals are exposed to increasing amounts of visual experience. We speculate that some aspects of variability in human visual behavior stem from these early system differences, which are broadly analogous to the stochasticity inherent in the random weight initialization of neural networks.

Our work breaks new ground by exploring human individual differences in visual behavior at the stimulus level. While individual differences in vision have been widely studied, this literature tends to focus on the overall task performance. For example, previous studies have documented individual differences in sensory encoding (Rahnev, 2021), color perception (Bosten, 2022), object recognition ability (Richler et al., 2019), etc. However, such task-level summaries provide only a coarse view of behavior: distinct individuals can have similar overall task performance but diverge substantially in their responses to specific stimuli. For example, two subjects may both achieve 70% accuracy on the same task while making errors on entirely different stimuli. In contrast to most existing work, we examined the individual differences in accuracy, confidence, and RT for each stimulus, revealing a high-dimensional structure of individual differences. These results demonstrate that individual differences in perceptual decision making extend far beyond task-level summaries, highlighting the importance of stimulus-level analysis and broadening the landscape of research on human perceptual variability.

In conclusion, we found that neural networks capture the structured individual variability in humans. Our results demonstrate that neural networks can serve as a proxy for studying individual differences in human behavior and highlight the need to develop models that are not only human-like on average but also human-like in their idiosyncrasy.

## 4 Methods

### 4.1 Human experiments

We reanalyzed data from an existing study where subjects performed an 8-choice noisy digit recognition task (Rafiei et al., 2024) and collected new data for a 10-choice blurry object recognition task.

#### 4.1.1 8-choice noisy digit recognition task (MNIST)

The dataset included 60 human subjects (Rafiei et al., 2024). Each trial began with the subject fixating on a white fixation cross for 500-1000 ms, followed by an image of a handwritten digit (between 1 – 8) embedded in low (0.25) or high (0.4) level of random noise, shown for 300 ms (Figure 1a). Subjects then sequentially reported their decision and confidence (4-point rating scale) with no time constraints. The stimulus set contains a total of 480 images randomly selected from the MNIST validation dataset (Deng, 2012), each tested twice for a total of 960 trials. On different trials, subjects were instructed to prioritize either speed (speed-focus) or accuracy (accuracy-focus). Responses to the two repetitions of each image were averaged in the analyses. More details for the experiment and data preprocessing procedures can be found in the original publication.

#### 4.1.2 10-choice blurry object recognition task (EcoSet10)

This study’s sample size, experimental design, included variables, hypotheses, and planned analyses were preregistered on the Open Science Framework (https://osf.io/n6m7b) prior to any data being collected.

As we preregistered, we targeted a total of 60 subjects after applying the exclusion criteria. A total of 64 subjects (38 females; age range = 18-26, mean = 19.4, SD = 1.46) were recruited from the student body at Georgia Tech. Per our preregistration, we excluded four subjects whose accuracy fell below 25% or deviated more than 2.5 standard deviations from the group mean. All subjects completed the experiment online and received 1 SONA credit as compensation. Subjects were naïve to the purpose of the study, reported normal or corrected-to-normal vision, and provided informed consent. Ethical approval for the study was obtained from the Georgia Tech Institutional Review Board (Protocol number: H21041).

The task stimuli consisted of 400 images from the EcoSet dataset (Mehrer et al., 2021). EcoSet is an ecologically motivated image dataset that incorporates linguistic corpus statistics and human ratings to define categories that are most relevant to humans. From the 565 categories in EcoSet, we selected those that met the following criteria: contains more than 4500 images in EcoSet, a concreteness rating above 4.75, and a frequency concreteness index greater than 0.5. From the resulting set, we then selected 10 categories to maximize perceptual and conceptual distinctiveness, avoiding closely related categories (e.g. dog and cat) to minimize confusion between categories. The final set had 10 categories: bed, bridge, car, cat, elephant, flower, house, knife, phone, and train. For each category, 40 unique images were randomly selected from the testing set of EcoSet to ensure the selected stimuli were novel for both human subjects and the trained models. All images were resized to 400×400 pixels for optimal viewing. Two levels of blurring were created by applying a Gaussian blur filter with a standard deviation of *σ*=6 and *σ*=10, using kernel sizes of 6*σ* + 1. The two blur levels were randomly interleaved throughout the experiment.

The experiment was programmed in JavaScript with the jsPsych library 7.3.3 (de Leeuw, 2015). The virtual chinrest method was used to control differences in viewing distance, display size, and stimulus size (Fung et al., 2025; Li et al., 2020). Specifically, subjects were first asked to resize an image of a credit card on the screen to match the size of a physical credit card, thus adjusting for discrepancies in display size. Then, subjects fixated on a central point and tracked the lateral movement of a target in their periphery, pressing the spacebar when the target disappeared (i.e., when it entered the blind spot). The procedure was repeated three times to estimate the viewing distance, assuming the blind spot is located at 13.5° of the visual angle.

Each trial began with a black fixation cross displayed for 300 ms, followed by a stimulus shown for 300 ms (Figure 5a). Subjects categorized the image by clicking one of the 10 category labels. They then rated their confidence on an 11-point scale from 0% to 100% in 10% increments. Both the perceptual and confidence responses had no time constraints, and the next trial began after both responses had been made.

Each of the 400 images was shown twice for a total of 800 trials, completed in the form of 4 runs, each containing 4 blocks of 50 trials. Before the main experiment, subjects completed two practice blocks. In the first block, subjects performed 10 trials with undistorted images to become familiar with the 10 categories and the task. This was followed by a second practice block of 20 trials where the images had the two levels of blur used in the experiment. Feedback was provided during the practice block but not during the main experiment. The full experimental procedure took about 60 minutes to complete.

We followed the same procedure as in the digit recognition task to remove individual trials with extreme RT values based on Tukey’s interquartile criterion (Rafiei et al., 2024). Specifically, for each subject and blur condition, we computed the 25^th^ (Q1) and 75^th^ (Q3) percentiles of the response time (RT) distribution. Trials with RTs outside the range [Q1 – 1.5 × (Q3 – Q1), [Q3 + 1.5 × (Q3 – Q1)] were excluded from further analyses. This procedure resulted in the removal of 8.76% of trials per subject (range = 2.75-13.8%). Note that this trial exclusion was not preregistered but was performed for consistency in the analyses of the two datasets.

### 4.2 General ANN training and testing procedures

We considered ANNs of three architectures: RTNet (Rafiei et al., 2024), AlexNet (Krizhevsky et al., 2012), and ResNet18 (He et al., 2015). For each architecture, we trained 60 instances to match the number of human subjects by varying only the random seed used for weight initialization. All model instances were trained on undistorted images from the dataset corresponding to the task (MNIST and EcoSet10). We then tested on the same images that were presented to the human subjects in the experiment and adjusted the stimulus difficulty (i.e., noise level in MNIST and blur level in EcoSet10) to align population-level accuracy between humans and models. For all instances, we analyzed the categorical decisions. We further computed the confidence of the ANNs as the logit difference between the first and second highest predicted class, a method shown to better predict human confidence than alternative approaches (Shekhar et al., 2025).

#### 4.2.1 8-choice noisy digit recognition task (MNIST)

We tested RTNet, AlexNet, and ResNet18 on the 8-choice noisy digit recognition task completed by the human subjects. All instances were trained on 60,000 noiseless MNIST training images. Prior to training, these images were resized to 227 × 227 pixels and normalized to have a mean of 0.1307 and a standard deviation of 0.3081. All models were trained to ceiling, achieving over 97% on the validation dataset with noiseless images. The last epoch of all instances was then tested on the 480 images presented to the human subjects.

#### 4.2.2 10-choice blurry object recognition task (EcoSet10)

We tested RTNet, AlexNet, and ResNet18 on the 10-choice blurry object recognition task completed by the human subjects. All instances were trained exclusively on the 10 categories tested in humans from EcoSet (EcoSet10; Mehrer et al., 2021). Prior to training, these images were resized to 227×227 pixels and normalized to the ImageNet mean (RGB means: 0.485, 0.456, 0.406) and standard deviation (RGB SDs: 0.229, 0.224, 0.225). All models were evaluated using images from the EcoSet validation dataset from the 10 trained categories. The last epoch of all instances was then tested on the 400 images presented to the human subjects.

### 4.3 ANN architectures

#### 4.3.1 RTNet

RTNet is a recently developed neural network specifically designed for capturing the human characteristics of perceptual decision-making (Rafiei et al., 2024). Unlike the other two standard feedforward convolutional networks, RTNet is a Bayesian neural network with probabilistic weights and incorporates an evidence accumulation mechanism, repeatedly processing an image until the accumulated evidence for a choice reaches a threshold. This mechanism allows RTNet to produce a model-equivalent of RT, defined as the number of repetition needed to reach the threshold. Additionally, RTNet generates stochastic decisions. Therefore, we tested the same images with ten repetitions and average their responses.

For the digit recognition task, we tested 60 RTNet instances that differ only in their random initialization during training. These instances were trained for 15 epochs with a batch size of 500, using the ELBO loss function and Adam optimizer with the default parameters (learning rate = 0.001; weight decay = 0, *ε* = 10^−8^). During testing, we matched the population-level accuracy between humans and models across the four experimental conditions (accuracy- vs. speed-focus, low- vs. high-noise) by searching stimulus noise levels from 0.2 to 12.0 in 100 evenly spaced steps, and the threshold level from 2 to 8 in increments of 0.5. The closest match to human accuracy was achieved for noise levels of 2.28 for low noise images and 4.25 for high noise images, and threshold values of 3 for the speed-focus condition and 6 for the accuracy-focus condition.

For the object recognition task, we trained 60 instances of RTNet that differ in their initializations. Unlike standard convolutional neural networks, RTNet weight initializations were performed by different combinations of means and standard deviations, allowing each weight distribution to capture distinct prior uncertainties. Specifically, the means of the weights and biases were set to a value between 0.1 and 1.2 with 0.1 increments, and all standard deviations of the weights and biases were set to a value ranging from 1 to 5 with increments of 1 (total: 12×5=60 instances). Models that failed to converge were retrained with the standard deviation sampled from 6 to 10 with increments of 1, ensuring that none of the instances has the same initialization distribution. We applied random horizontal flipping (50%) to the image input during training to reduce overfitting and improve model generalization. All instances were trained for 150 epochs with a batch size of 512, using the ELBO loss function and Adam optimizer with learning rate = 10^−5^; weight decay = 0, *ε* = 10^−8^. The KL divergence term in the ELBO loss function was gradually scaled using KL annealing, with the KL weight *β* following a sigmoid schedule:

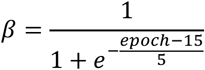

The gradually increasing KL divergence term enables the model to first focus on fitting the data in early training and then progressively regularize the weights according to the priors. All instances achieved above 75% accuracy in the validation dataset. During testing, we matched the population-level accuracy between humans and models across two blur conditions by adjusting the blur level (search from 0.5 to 2 in increments of 0.025). The closest match to human accuracy was achieved by *σ* = 0.9 and *σ* = 1.525 for low and high blur, respectively.

#### 4.3.2 AlexNet

AlexNet is a standard feedforward convolutional neural network with five convolutional layers and three fully connected layers (Krizhevsky et al., 2012). For both tasks, we initialized 60 different random weight initializations using Kaiming uniform initialization and trained them independently as detailed below.

For the digit recognition task, all instances were trained for 15 epochs with a batch size of 128, using the cross-entropy loss function and the Adam optimizer with default parameters (learning rate = 0.001; weight decay = 0, *ε* = 10^−8^). During testing, since AlexNet has no inherent adjustment for threshold, we matched the population-level accuracy between humans and models across the four experimental conditions only by adjusting the noise level (search from 1 to 3.5 in increments of 0.01). The closest match to human accuracy was achieved by 1.425 and 2.7 for low and high noise in the accuracy-focus condition, and 1.75 and 2.825 for low and high noise in the speed-focus condition.

For the object recognition task, we applied random horizontal flipping (50%) to the image input during training. All instances were trained for 30 epochs with a batch size of 512, using the cross-entropy loss function and the Adam optimizer with learning rate = 10^−5^; weight decay = 10^−4^ and *ε* = 10^−8^. The learning rate was adjusted using a reduce-on-plateau scheduler that monitored validation loss and reduced the learning rate by a factor of 0.5 when no improvement was observed for three consecutive epochs. All instances achieved above 80% accuracy in the validation dataset. During testing, we matched the population-level accuracy between humans and models across two blur conditions by adjusting the blur level (search from 0.5 to 2 in increments of 0.025). The closest match to human accuracy was achieved by *σ* = 1.25 and *σ* = 1.90 for low and high blur, respectively.

#### 4.3.3 ResNet18

ResNet18 is a convolutional architecture with residual connections that incorporate skip connections, allowing information to bypass intermediate layers for training deeper networks (He et al., 2015). For both tasks, we initialized 60 different random weight initializations using Kaiming uniform initialization and trained them independently as detailed below.

For the digit recognition task, all instances were trained for 15 epochs with a batch size of 128, using the cross-entropy loss function and the Adam optimizer with default parameters (learning rate = 0.001; weight decay = 0, *ε* = 10^−8^). During testing, we matched the population-level accuracy between humans and models across the four experimental conditions only by adjusting the noise level (search from 0 to 5 in increments of 0.1). The closest match to human accuracy was achieved by 0.52 and 0.86 for low and high noise in the accuracy-focus condition, and 0.6 and 0.9 for low and high noise in the speed-focus condition.

For the object recognition task, we applied random horizontal flipping (50%) and random rotation (15°) to the image input during training. All instances were trained for 30 epochs with a batch size of 512, using the cross-entropy loss function and the Adam optimizer with learning rate = 10^−5^; weight decay = 10^−4^ and *ε* = 10^−8^. The learning rate was adjusted using a reduce-on-plateau scheduler that monitored validation loss and reduced the learning rate by a factor of 0.5 when no improvement was observed for two consecutive epochs. All instances achieved above 85% accuracy in the validation dataset. During testing, we matched the population-level accuracy between humans and models across two blur conditions by adjusting the blur level (search from 0.5 to 2 in increments of 0.025). The closest match to human accuracy was achieved by *σ* = 1.225 and *σ* = 1.725 for low and high blur, respectively.

### 4.4 Analyses

#### 4.4.1 Testing the alignment of individual ANN instances to individual human subjects

To test the alignment of individual ANN instances to individual human subjects, we first organized the data into the form of a response matrix *R*, where each row represents an individual (human or ANN instance) and each column represents an image:

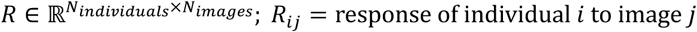

For human subjects and RTNet instances (which generate stochastic decisions), we averaged their responses across the two stimulus repetitions in the response matrix. In contrast, AlexNet and ResNet18 produce deterministic outputs for a given image, so no averaging was performed for these models. Thus, for both humans and different ANN architectures, we created a response matrix for each behavioral metric (accuracy, confidence, and RT).

We then computed the pairwise correlations between the responses of each human subject and each ANN instance to obtain a similarity matrix (*S*) of size *N*_ℎ*uman*_ × *N_ANN_*, where each entry is computed as the dot product of *z*-scored response matrix, which is mathematically equivalent to the Pearson correlation. Higher correlation values indicate better alignment between a human subject and an ANN instance:

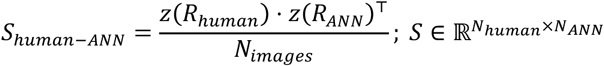

Note that to compute the similarity matrix, *S*, we used all images regardless of their difficulty level. Further, the 8-choice digit recognition task included speed and accuracy-focused conditions. Since these can be seen as slightly different tasks, we computed the similarity matrix separately for each of these two conditions and then averaged them to obtain a single similarity matrix.

For benchmarking, we also computed human-human similarity matrices using the same procedure. Since the diagonal of the human-human similarity matrix is always 1 (as it corresponds to an individual being correlated with themselves), we removed the diagonal in subsequent analyses.

#### 4.4.2 Statistical properties of the human-human and human-ANN similarity matrices

We first examined the basic statistical properties of the human-ANN and human-human similarity matrices to gain an overall understanding of how individual human subjects align with ANN instances and with one another. We computed the standard deviation of the similarity matrix to quantify variability. To enable statistical comparison between the variability of the human-ANN and human-human similarity matrices, we performed 1000 iterations of randomly splitting the images into two equal halves, yielding a total of 2000 subsets. For each subset, we computed the corresponding similarity matrix and its standard deviation. We then assessed statistical significance using a non-parametric bootstrap test. Specifically, for each bootstrap sample, we computed the difference in standard deviation between human-ANNs and human-human similarity matrix. The two-sided p-value was the proportion of bootstrap samples where the difference crossed zero, and the 2.5^th^ and 97.5^th^ percentiles provided a 95% confidence interval. For completeness, we also examined mean similarity. For each subject, we computed the average correlation with all instances or all other subjects (excluding the target subject). Differences in mean similarity between human-ANN and human-human were tested using paired-sample t-tests.

#### 4.4.3 Consistency of mapping between ANN instances and human subjects

To ensure that the mapping between individual ANN instances and individual human subjects is not spurious, we tested the consistency of this mapping across different subsets of the data. Specifically, we randomly split the images into two equal halves and computed the corresponding similarity matrix for each split. This allowed us to quantify the consistency of the resulting mappings across the two halves of the data. This procedure was repeated for 1000 bootstrap iterations with different random image splits to ensure our results were not specific to a particular data subset.

Consistency between the two similarity matrices was quantified in two ways: (1) by Pearson correlation, and (2) by rank. First, for Pearson correlation, we computed the correlation between the two splits within each individual subject. We then corrected this value by subtracting the average correlation between the same subject and all other subjects, which removes any correlations stemming from ANN instances that may perform uniformly well or uniformly badly in matching all human subjects. Mathematically, this is defined as:

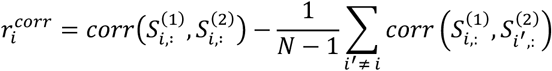

where 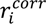 refers to the corrected Pearson correlation for subject *i*; 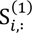 and 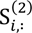 are the rows corresponding to subject *i* in the similarity matrices from splits 1 and 2, respectively; *N* is the total number of subjects; and the summation runs over all other subjects without self-correlations, *i*^′^ ≠ *i*.

The corrected correlation consistency values for each subject were then averaged across 1000 bootstrap iterations. We statistically compared the human-human and human-ANN correlation consistencies using paired t-test across subjects. In addition, we computed the Bayes factor for this comparison using the default Cauchy scale factor in the Pingouin Python package (https://pingouin-stats.org/build/html/index.html).

Since using Pearson correlation in measuring consistency could partially be driven by a few ANN instances that always show good or bad alignment with most subjects, we conducted a complementary analysis by quantifying consistency using the rank order relationship. Specifically, for each subject, we ranked all ANN instances (or each subject except the target subject) based on their degree of alignment with the target subject. For each subject *i* and each unique pair of ANN instances (or subjects) (*j*_1_, *j*_2_) with *j*_1_ < *j*_2_, we computed the rank differences separately in each split:

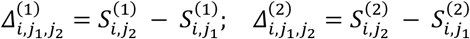

where *S*^(*k*)^ denotes the ranked similarity between subject *i* and ANN instance (or subject) *j* in split *k*. These differences Δ*_i_*_,j0_,j_1_ were then multiplied across the two splits:

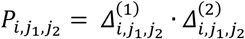

The idea behind the multiplication is that positive products would indicate that the relative ordering of the pair of ANN instances (or subjects) is consistent across splits, which would increase the score multiplicatively, whereas negative products indicate the ordering flips between splits and are penalized. Finally, we averaged over all pairs and all subjects to obtain a single rank consistency score:

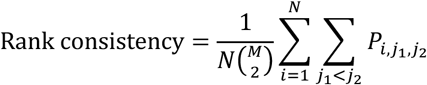

where *N* is the total number of subjects and *M* is the total number of items being ranked: *M* = *N* for the human-ANN mapping, and *M* = *N* − 1 for the human-human mapping.

To assess statistical significance of the differences between human-human and human-ANN mappings in the rank consistency metric, we used a non-parametric bootstrap test. Specifically, for each of 1000 bootstrap iteration, we compared the difference in the rank consistency score between human-human and human-ANN similarity matrices. The proportion of bootstrap samples with differences greater than or less than zero was used to calculate a two-sided *p*-value, and the 2.5^th^ and 97.5^th^ percentiles of the bootstrap differences were used to obtain a 95% confidence interval.

#### 4.4.4 Across-metric analyses

To investigate whether the mapping consistency between individual ANN instances and individual human subjects extends across the three behavioral metrics (accuracy, confidence, and RT), we applied a procedure analogous to that described above, but computed correlations between similarity matrices obtained from pairs of behavioral metrics (e.g. accuracy vs. confidence). Statistical significance was assessed by comparing human-human and human-ANN correlations against zero using one-sample t-tests. We then performed the same mapping consistency analyses (corrected Pearson correlation and rank consistency) to confirm that these across-metric mappings reflect systematic individual differences. Critically, we evaluated the across-metric consistency by comparing the first split of data of one metric with the second split of the other metric, ensuring that the measured consistency was not driven by carryover effects within the same subset of images.

#### 4.4.5 Mapping analyses across different levels of average performance

Under the current evaluation, we adjusted the noise level of the stimulus to calibrate the overall performance of the ANN population to that of human subjects. To examine how the mapping between humans and ANNs varied across neural network populations spanning different levels of performance, for each ANN architecture, we used five levels of stimulus noise so that the average accuracy of the ANN population was equal to *mean* − 2*SD*, *mean* − 1*SD*, *mean*, *mean* + 1*SD*, and *mean* + 2*SD* of the human performance distribution, which were 51.8%, 61.6 %, 71.4 %, 81.2%, and 91.0%, respectively. Specifically, for each ANN architecture, we searched through a range of stimulus noise levels and found the noise levels that matched the human accuracy across the four experimental conditions. Note that for each architecture, the ANN population that corresponded to the human mean is the result that we reported in the main paper (Figure 2d for within-metric & Figure 3c for across-metric). The above procedure produces a total of 15 ANN populations (3 ANN architectures x 5 levels of performance). For each of the ANN population, we repeated the same mapping consistency analyses with the human data as described above. We then computed the statistical significance of the human-ANN mapping strength against zero using one-sample t-tests and against the human-human benchmark using paired t-tests.

#### 4.4.6 Pseudo-instances analyses

To determine whether individual human-ANN mappings capture more than overall performance similarities, we have to dissociate instance-specific differences in ANNs from differences in overall performance. Thus, we created “pseudo-instances” from a single ANN instance by systematically varying the level of stimulus noise during testing. For each ANN instance, we generated 60 pseudo-instances. This allowed us to produce a range of performance levels across pseudo-instances while keeping the underlying ANN instance identical. We then constructed the mapping between 60 human subjects and 60 pseudo-instances and performed the same within-metric and across-metric mapping consistency analyses as described before. Since there are 60 true instances for each architecture, we repeated this procedure for each of the 60 true instances and averaged their results. We compared these results with the mapping consistency results using the 60 true instances that we reported in the main paper (Figure 2d for within-metric & Figure 3c for across-metric). Statistical significance was assessed by comparing the differences between the mapping consistency with pseudo-instances and true instances using paired t-tests.

#### 4.4.7 Untrained network analyses

To understand whether neural network training contributed to the mapping between individual human subjects and ANN instances, we tested the alignment between 60 human subjects and 60 untrained RTNet instances. Specifically, we took the same 60 initializations of RTNet that we used in the paper and tested their performance on the 480 test images prior to training. The noise level and response threshold were set to the values at which the corresponding trained RTNet population achieved the closest match to human accuracy (Noise level: low noise: 2.1, high noise: 4.1; Response threshold: speed focus: 3, accuracy focus = 6). We then performed the same mapping procedure to construct the similarity matrices between 60 human subjects and 60 untrained RTNet instances and assessed their mapping consistency.

We compared the mapping results from the untrained RTNet instances with the trained RTNet instances that we reported in the main paper (Figure 2d for within-metric & Figure 3c for across-metric). Statistical significance was evaluated using paired t-tests between the untrained and trained instances. We did not include untrained AlexNet and ResNet18 in this figure because they produced the same behavioral response for all or almost all images, preventing the computation of image-by-image correlations required for the mapping analysis.

#### 4.4.8 Alignment-weighted prediction analysis

To test whether human-ANN mappings could improve behavioral predictions for unseen data, we performed a cross-data subset prediction analysis. Specifically, we computed the human-ANN similarity matrix from one data subset to learn the alignment between each subject and ANN instance. These correlations in the similarity matrix were then used as weights to combine ANN responses, predicting the human response matrix in the held-out data subset. Mathematically:

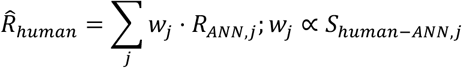

where *j* refers to the index of instance, *R*°_ℎ*uman*_refers to the response of ANN instance on the held-out data subset, and *w*_j_ is proportional to the human-ANN similarity matrix estimated from the training subset.

The above procedure was repeated for 1000 bootstrap iterations with different random image splits to ensure our results were not specific to a particular data subset. To assess the benefit of the alignment weighting, we compared the weighted prediction with an unweighted average prediction that simply averaged responses across all ANN instances. We then evaluated the statistical significance of the improvement by computing the difference between the alignment-weighted and unweighted predictions and testing the difference against zero using one-sample t-tests. Additionally, these differences were compared to the human-human mapping benchmark using paired-sample t-tests.

#### 4.4.9 Digital twin analysis

We predicted the behavioral responses of individual human subjects using a digital twin approach. Specifically, for each behavioral metric, we computed the human-ANN similarity matrix from one data subset to identify the best matching ANN instance for a specific human subject (a.k.a. digital twin). In a held-out data subset, we computed the image-by-image correlation between the subject’s responses and digital twin’s responses (prediction) to evaluate prediction accuracy of the digital twin. We then compared the digital twin prediction accuracy with the image-by-image correlations between the subject’s response and each of the remaining instances and averaged across all other instances. To create a human benchmark, we followed the same procedure but instead we identified the best-matching other subject that best aligns with the target subject and compared the best-matching subject’s prediction with all other subjects.

The above procedure was repeated for 1000 bootstrap iterations with different random image splits and the results were averaged across iterations. To assess statistical significance of the prediction benefit from the digital twin/best-matching other subject, we computed the difference between the digital twin’s prediction and all other instances’ predictions and compared the difference against zero using one-sample t-tests. Additionally, comparison with the human benchmark was done using paired-sample t-tests.

### 4.5 Data and code availability

All data and codes are publicly available at https://github.com/herrickfung/midb_data_code.

The full analysis pipeline is packaged separately in a custom Python library, *IndiMap*, for analyzing and mapping individual differences between humans and artificial neural networks. The full repository is available at https://github.com/herrickfung/indimap.

## Author Contributions

All authors jointly designed the research, interpreted the results, and contributed to revising and editing the manuscript. H.F. conducted the research, collected and analyzed the data, and drafted the manuscript. N.A.R.M. and D.R. reviewed the manuscript.

## Acknowledgment

This work was supported by the National Institute of Health through grant award R01MH119189 to D.R. and R00EY032603 to N.A.R.M. We thank members of the Rahnev and Murty labs for thoughtful discussions.

## 6 Supplementary Results

### 6.1 Wasserstein distance in humans and ANN instances

In the main text, we demonstrated substantial individual differences in humans and ANN instances based on summary statistics computed for each individual (Figure 1b). However, these summary statistics do not capture differences at the level of trial-by-trial response distributions. To directly assess distributional differences in humans and ANNs, we computed the pairwise Wasserstein distance between the trial-level response distributions for every possible pair of individual human subjects or individual ANN instances (*C*^60^ = 1770). Formally, for individuals *i* and *j*, the Wasserstein distance was defined as:

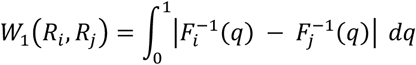

where *R_i_* and *R*_j_ denote the trial-by-trial response from individual *i* and *j*, respectively, and *F*^−1^(*q*) and

*F*^−1^(*q*) are the quantile functions (inverse cumulative distribution functions), giving the values at the *q*-th percentile of each individual’s trial responses. In other words, the Wasserstein distance measures the average difference between two individual’s response across all percentiles, with larger values indicating larger variability.

For the 8-choice digit recognition task, we found that RTNet and AlexNet generally showed slightly greater individual variability than humans for accuracy (Human: 0.091 ± 0.076; RTNet: 0.141 ± 0.098; AlexNet: 0.177 ± 0.143; Supplementary Figure 1) and confidence (Human: 0.150 ± 0.112; RTNet: 0.398 ± 0.313; AlexNet: 0.424 ± 0.305), whereas ResNet18 exhibited a similar level of variability (Accuracy: 0.101 ± 0.077; Confidence: 0.138 ± 0.078). For RT, however, RTNet showed slightly smaller variability than humans (Human: 0.170 ± 0.112; RTNet: 0.130 ± 0.089). Overall, these results replicate the results from Figure 1b by demonstrating that ANNs that only differ in their random initialization exhibit substantial individual differences that either match or exceed the variability among human subjects.

For the 10-choice object recognition task, ANN instances generally showed markedly reduced variability as compared to humans for both accuracy (Human: 0.104 ± 0.093; RTNet: 0.040 ± 0.028; AlexNet: 0.020 ± 0.014; ResNet18: 0.029 ± 0.021; Supplementary Figure 8) and RT (Human: 0.225 ± 0.157; RTNet: 0.105 ± 0.119). The only exception was confidence, for which RTNet demonstrated higher variability than humans (Human: 0.264 ± 0.156; RTNet: 0.375 ± 0.316), while AlexNet (0.062 ± 0.025) and ResNet18 (0.080 ± 0.040) showed substantially reduced variability. These results showed that ANNs had substantially less variability than the human subjects in the 10-choice object recognition task.

### 6.2 Comparing distributions of best-matching subjects and instances

The similarity matrices in Figure 2a and Supplementary Figure 2 demonstrate that the same human subject or same ANN instance sometimes provides the best match to the data for multiple human subjects. If individual differences in ANN instances are a good model of individual differences in human subjects, we may expect that this effect would be comparable in human-human and human-ANN similarity matrices.

To investigate whether a small subset of subjects or ANN instances dominated the alignment with most human subjects, we analyzed the distribution of best-match instances (or subjects) across repeated splits of the data. Specifically, we performed 1000 iterations of randomly splitting the images into two equal halves, yielding a total of 2000 subsets. For each subset, we computed a similarity matrix and identified, for each subject, the ANN instance (or subject) which has the highest similarity. We then counted how often each ANN instance (or subject) was selected as the best match across all human subjects. These counts were ranked in descending order to obtain the best-match frequency distribution (Supplementary Figure 3a), which we modeled with an exponential decay function:

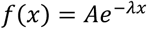

where *A* is a scaling factor, and λ is the decay rate, indicating how quickly best-match frequency decreases down the ranked list of instances/subjects. Comparing the decay rates between humans and different ANN architectures allowed us to quantify differences in their alignment structure. Statistical significance in the decay rates was assessed using a non-parametric bootstrap test. For each bootstrap sample, we computed the difference in the decay rate between humans and ANNs. The two-sided p-value was the proportion of bootstrap samples where the difference crossed zero, and the 2.5th and 97.5th percentiles provided a 95% confidence interval.

We found that the best-match frequency distributions for humans and ANN instances had very similar decay rates (Supplementary Figure 3b). Specifically, the decay rate for humans was not statistically different from that of any of the three network architectures for any metric (all *p*’s > 0.129). These results demonstrate that the best-match frequency distributions, which describe the variability in the similarity of individual subjects or ANN instances to the 60 human subjects, are broadly comparable for humans and ANN instances.

## 7 Supplementary Figures

**Supplementary Figure 1.**
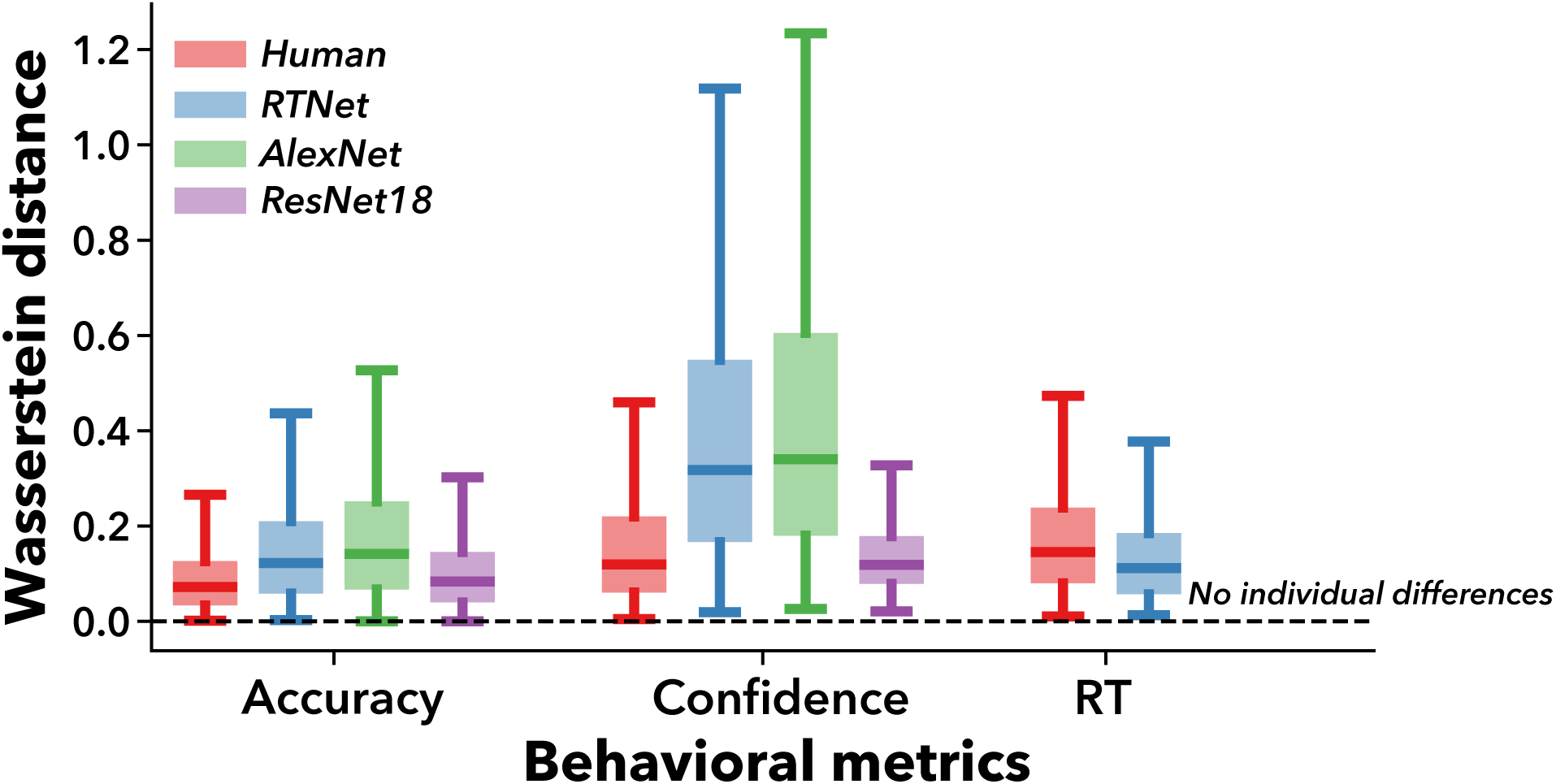
Wasserstein distance for humans and ANN instances for the 8-choice digit recognition task. Figure 1b demonstrated substantial individual differences in humans and ANN instances based on summary statistics computed for each individual. However, these summary statistics do not capture differences at the level of trial-by-trial response distributions. To directly assess distributional differences, we computed the Wasserstein distances between the trial-level response distributions for each pair of human subjects and for each pair of individual ANN instances, separately for the three ANN architectures. Larger distance indicates larger individual variability. Consistent with the main text, both humans and ANNs exhibited substantial individual differences. RTNet and AlexNet generally showed slightly greater individual variability than humans for accuracy and confidence, whereas ResNet18 exhibited a comparable level of variability. For RT, RTNet showed slightly lower variability than that observed in humans. Boxes indicate the 25^th^ and 75^th^ percentiles, the central line denotes the median, and whiskers extend to 1.5 x IQR. The dashed line at zero indicates the baseline with no individual differences.

**Supplementary Figure 2.**
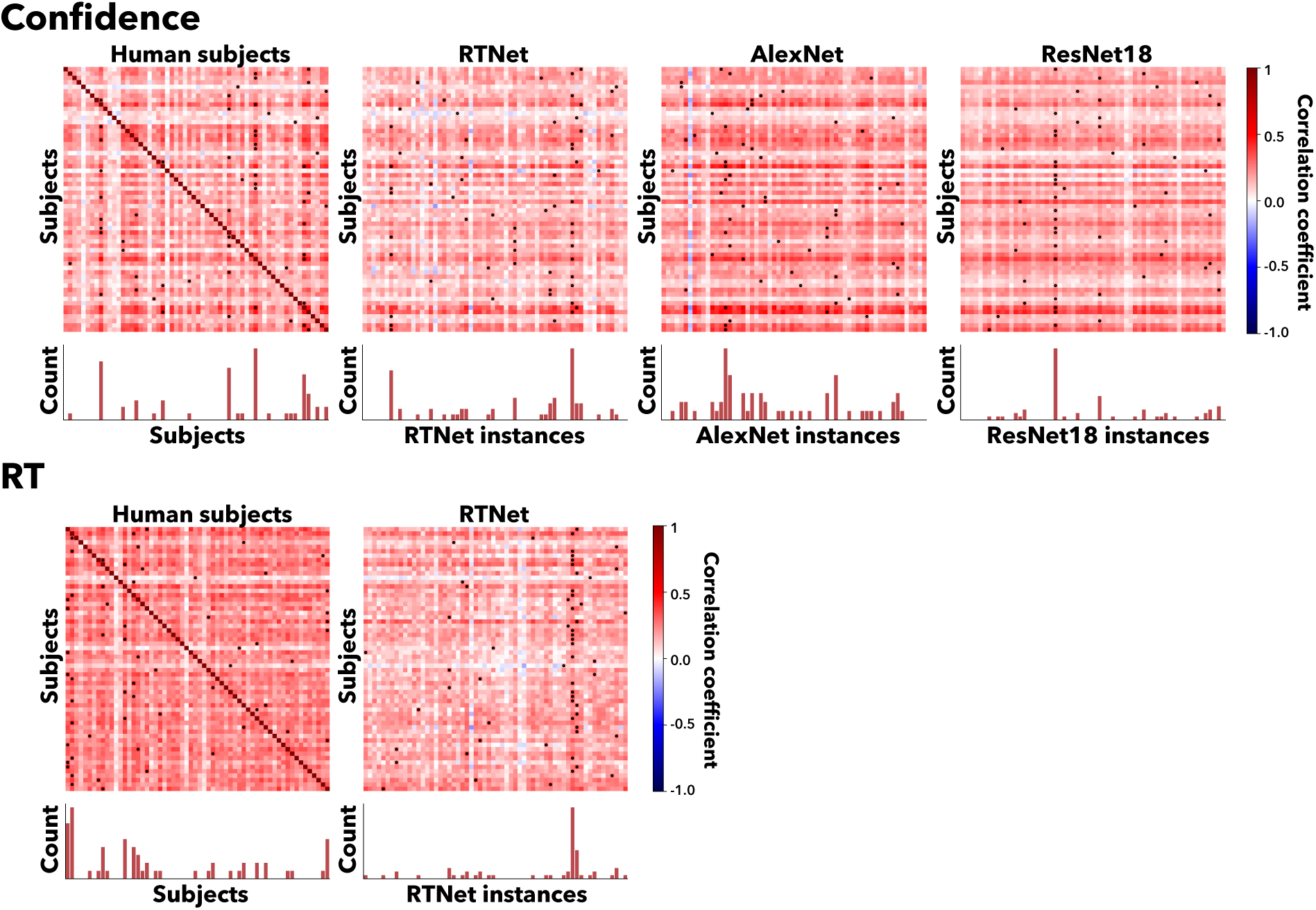
Similarity between each subject and each ANN instance in confidence and RT. We constructed the similarity matrix of 60 human subjects and the 60 instances of each ANN architecture, with higher values indicating a better alignment between an individual human and a specific ANN instance. The black dots indicate the model instance with the best fit for each subject (i.e., there is one black dot for each row). The histograms underneath show how often each subject or ANN instance was the best match across all human subjects. Consistent with the similarity matrix for accuracy as reported in Figure 2a, we observed a wide range of correlation and variability from aligning individual ANN instances to individual human subjects across three ANN architectures for confidence and RT. We also observed variability in the model instances that provided the best fit for each subject. We further explored these differences in variability in the main text and in Supplementary Figure 3.

**Supplementary Figure 3.**
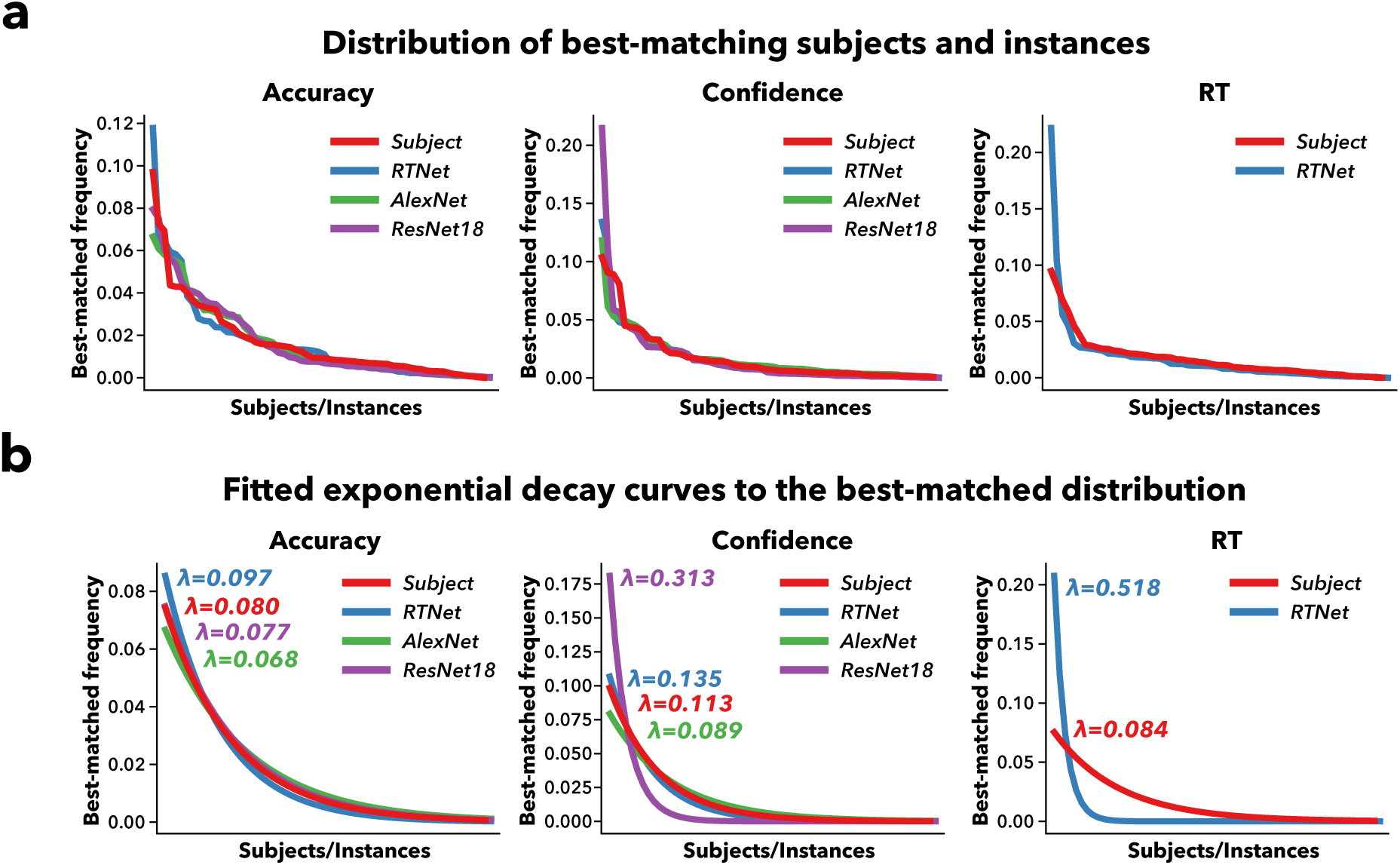
Distribution of best-matching subjects and instances and its exponential decay curve fitting. (a) Distribution of best-matching subjects and instances. The figure shows the frequency of each instance (or subject) being identified as the best match in descending order, where we observed that human and ANN instances have a relatively similar rate of decay. (b) The fitted exponential decay curve to the best-matched distribution. **λ** denotes the decay rate. The similarity matrices in Figure 2a and Supplementary Figure 2 show that the same human subject or same ANN instance sometimes provides the best-match to the data for multiple human subjects. To quantify and compare this effect between the human-human and human-ANN similarity matrices, we counted how often each subject or ANN instance was the best match across all human subjects. We then sorted these counts in descending order to obtain “best-match” frequency distribution. We modeled these distributions using an exponential decay function, allowing us to compare the decay rate between humans and ANNs. We performed this procedure in 2000 data subsets for statistical comparison. For all three behavioral metrics, we found that the decay rate for humans was not statistically different from RTNet, AlexNet, and ResNet18 (all *p*’s > 0.129). These results demonstrate that the best-match frequency distributions, which describe the variability in the similarity of individual subjects or ANN instances to the 60 human subjects, are broadly similar for humans and ANN instances.

**Supplementary Figure 4.**
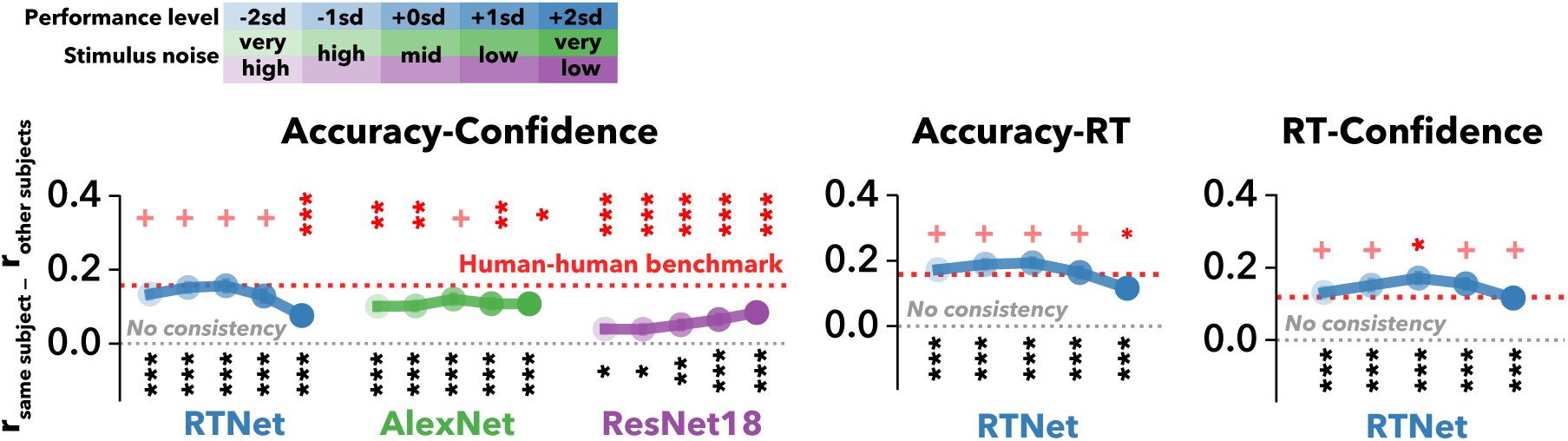
Human-ANN across-metric mapping consistency across five levels of average performance. To investigate how ANN populations that vary in their overall performance affect the individual human-ANN mapping, we used five levels of stimulus noise so that average accuracy of the ANN population was equal to *mean* − 2*SD*, *mean* − 1*SD*, *mean*, *mean* + 1*SD*, and *mean* + 2*SD* of the human performance distribution. We then repeated the same mapping analyses for each population. Similar to what we found using within-metric mapping consistency in Figure 4a, the strengths of the human-ANN mappings were largely similar across the three highest-noise stimuli. Yet, the two lowest stimulus noise levels demonstrated lower mapping consistency for all three pairs of behavioral metrics, which was likely due to a ceiling effect. Overall, these results demonstrate that the human-ANN across-metric mappings are generally consistent across different levels of overall ANN performance and do not only appear when performance is matched across humans and ANNs. Note that the mapping results of standard human-average population correspond to the results reported in the main text (0 SD; & Figure 3c for across-metric). Grey and red dotted lines denote the absence of mapping consistency and the human-human mapping benchmark, respectively. Black asterisks indicate significance against zero. Red asterisks indicate pairwise-comparisons with the human-human benchmark. For both, *, *p* < 0.05; **, *p* < 0.01; ***, *p* < 0.001, +, *p* > 0.05.

**Supplementary Figure 5.**
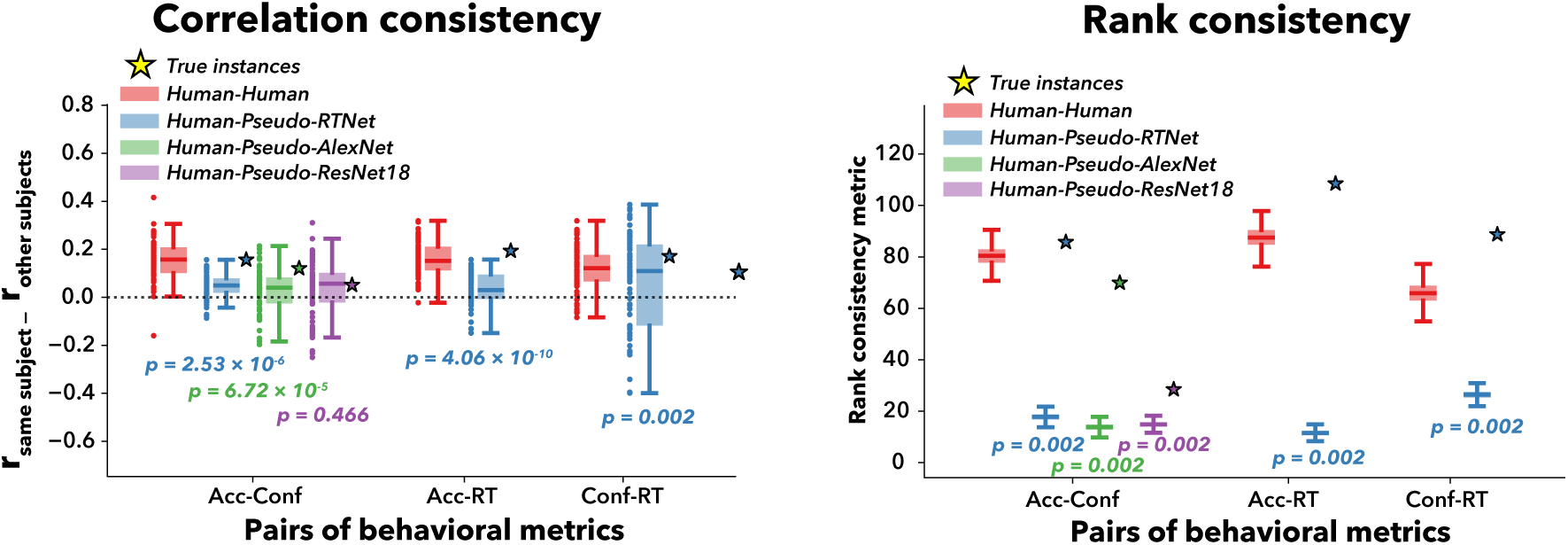
Human-ANN across-metric mapping consistency in pseudo-instances. To demonstrate that human-ANN mappings do not simply reflect similarities in overall performance, we created “pseudo-instances” from a single ANN instance by systematically varying the level of stimulus noise during testing. We then repeated the same mapping analyses on these pseudo-instances and compare these results with the mapping analyses on the true instances. Similar to what we observed for within-metric mapping consistency (Figure 4b), we found that the true instances showed significantly higher mapping consistency than their pseudo-instances counterparts in RTNet for all three pairs of behavioral metrics and for both the correlation and rank analyses. AlexNet also showed greater mapping consistency in true instances than pseudo-instances for accuracy-confidence in both correlation and rank analyses. ResNet18, however, showed greater mapping consistency for accuracy-confidence only for the rank analyses but not the correlation analyses. These results indicate that the across-metric mapping between individual differences in humans and ANN instances is driven by instance-specific factors beyond similarities in overall performance. Also note that the mapping consistencies for all pseudo- instances were significantly larger than zero across all behavioral metrics and ANN architectures, which shows that similarities in overall performance do contribute to the human-ANN mapping. However, the lower consistency in pseudo-instances than true instances demonstrates that performance similarity alone cannot fully account for the correspondence we observed between individual human subjects and true ANN instances. P-values indicate the significance of pairwise differences between the true instances (Star) and pseudo-instances.

**Supplementary Figure 6.**
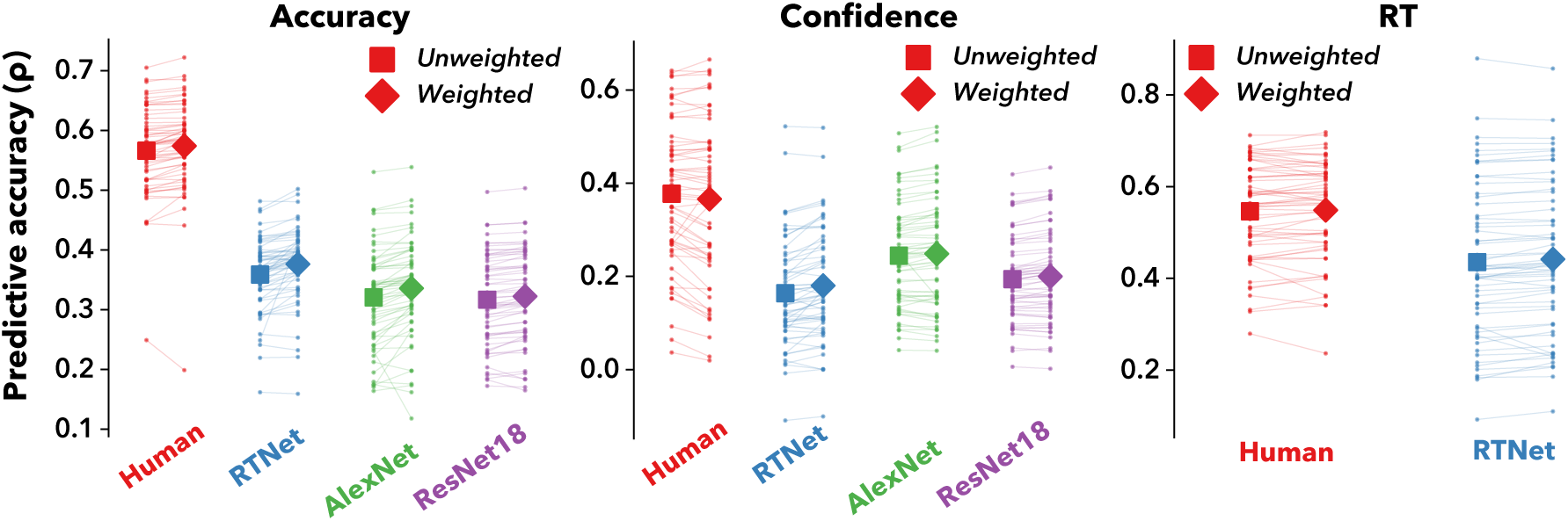
Raw prediction accuracy for alignment-weighted vs. unweighted prediction for the 8-choice digit recognition task. We used the correlations in the similarity matrix as weights to predict behavior in a held-out data subset. For comparison, we computed an unweighted prediction that simply averaged across all subjects or ANN instances. Across all three behavioral metrics, human subjects consistently exhibited higher prediction accuracy than ANNs, regardless of the weighting strategy. For all ANN architectures, the alignment-weighted predictions showed statistically significant improvements over the unweighted control. Although these improvements were modest in magnitude, they were highly consistent across different subjects. We analyzed the differences in improvement magnitude across humans and three ANNs in Figure 4c of the main text. Dots show individual human subjects, lines connect the alignment-weighted and unweighted predictions for the same subjects.

**Supplementary Figure 7.**
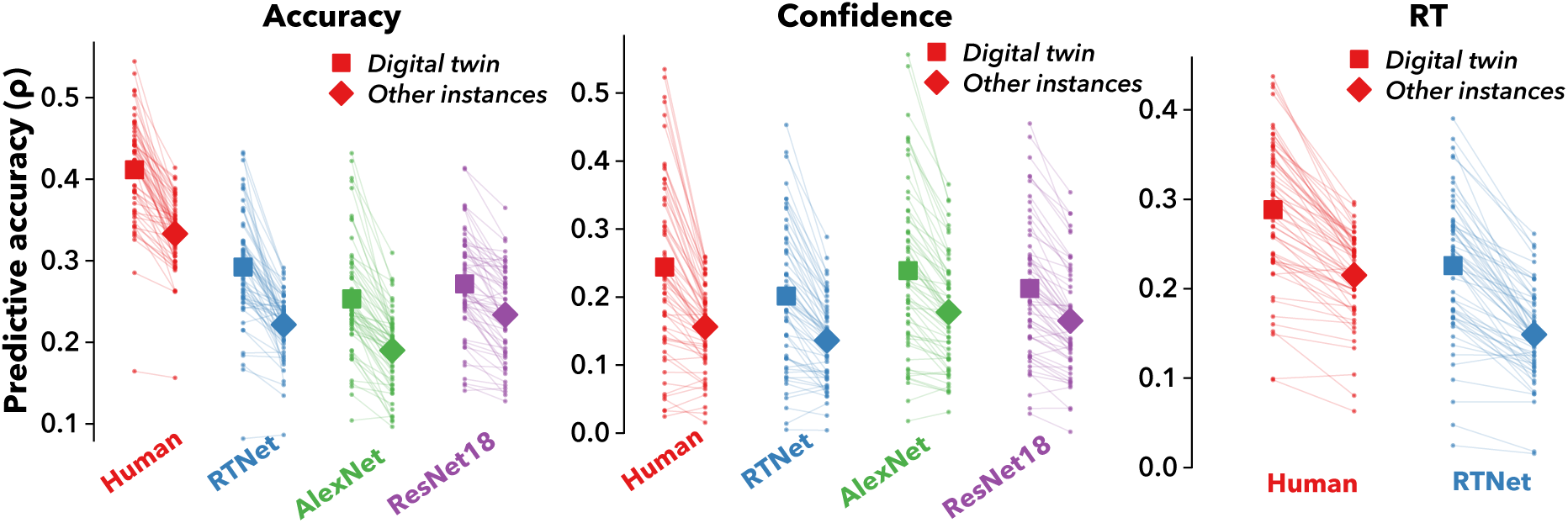
Raw prediction accuracy for predicting single-subject responses with the digital twin vs. all other instances for the 8-choice digit recognition task. We identified the best-matching subject-instance pair (digital twin). We then predicted each subject’s behavior in a held-out data subset using only the responses from the digital twin. For comparison, we also predicted behavior using the responses from other instances. As before, we see that human subjects generally exhibited higher prediction accuracy than ANNs for accuracy, but showed a similar level of prediction accuracy for confidence and RT. We analyzed the differences in improvement magnitude between the digital twin and the remaining instances in Figure 4d of the main text. Dots show individual human subjects, lines connect the digital twin predictions and other instances predictions for the same subjects.

**Supplementary Figure 8.**
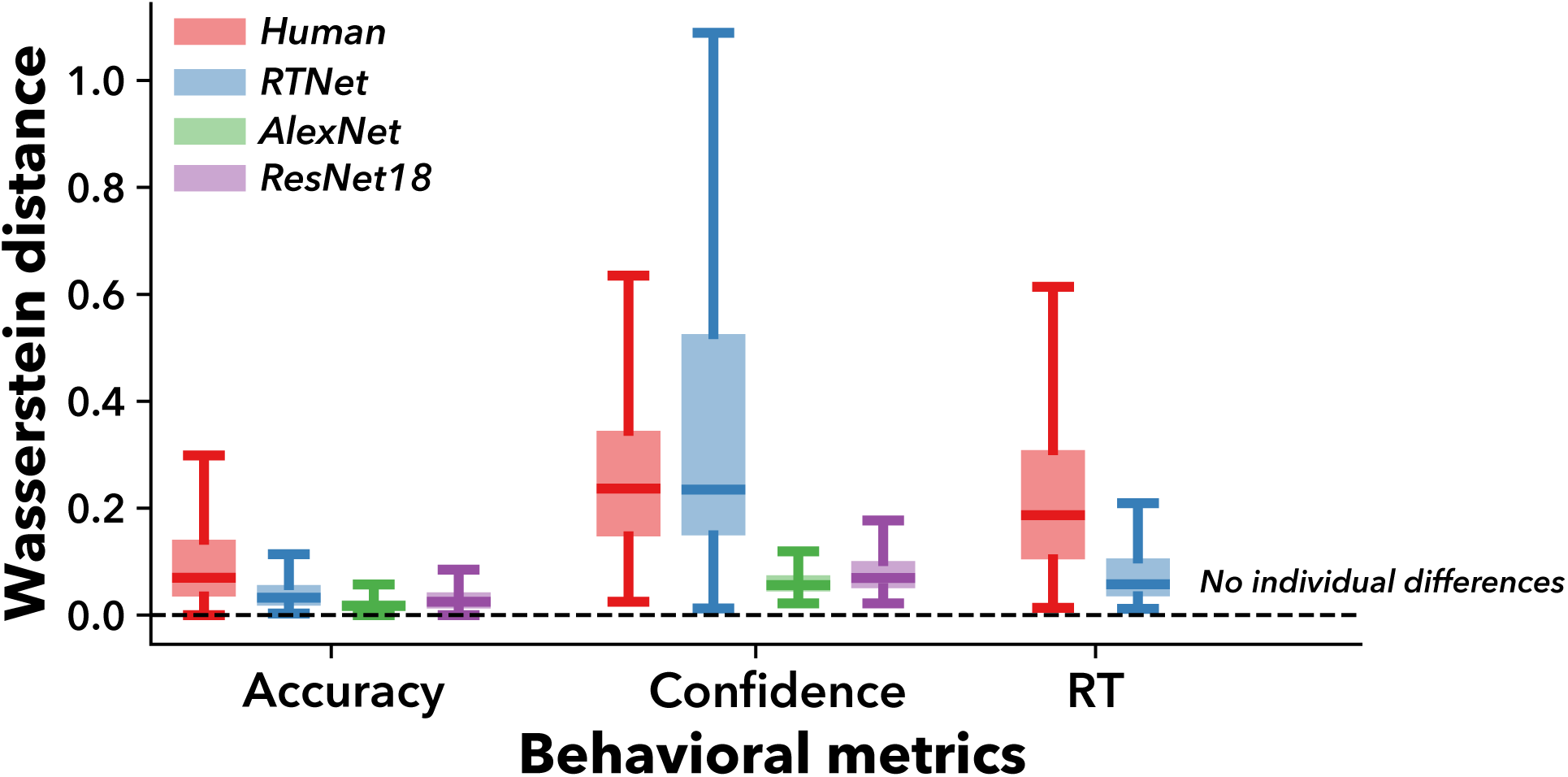
Wasserstein distance in humans and ANN instances for the 10-choice object recognition task. Figure 5b demonstrated the individual differences in humans and ANN instances using summary statistics computed for each individual. However, these summary statistics do not capture differences at the level of trial-by-trial response distributions. To directly assess distributional differences, we computed the pairwise Wasserstein distance between the trial-level response distributions for each pair of human subjects and for each pair of individual ANN instances, separately for the three ANN architectures. Larger distance indicates larger individual variability. Consistent with the main text, we found that ANN instances showed markedly reduced variability as compared to humans for both accuracy and RT. The only exception was confidence, for which RTNet demonstrated higher variability than humans, while AlexNet and ResNet18 showed substantially reduced variability. These results showed that ANNs had substantially less variability than the human subjects in the 10-choice object recognition task. Error bars show standard deviation across all subject/instance pairs. Dashed line at zero indicates the baseline with no individual differences.

**Supplementary Figure 9.**
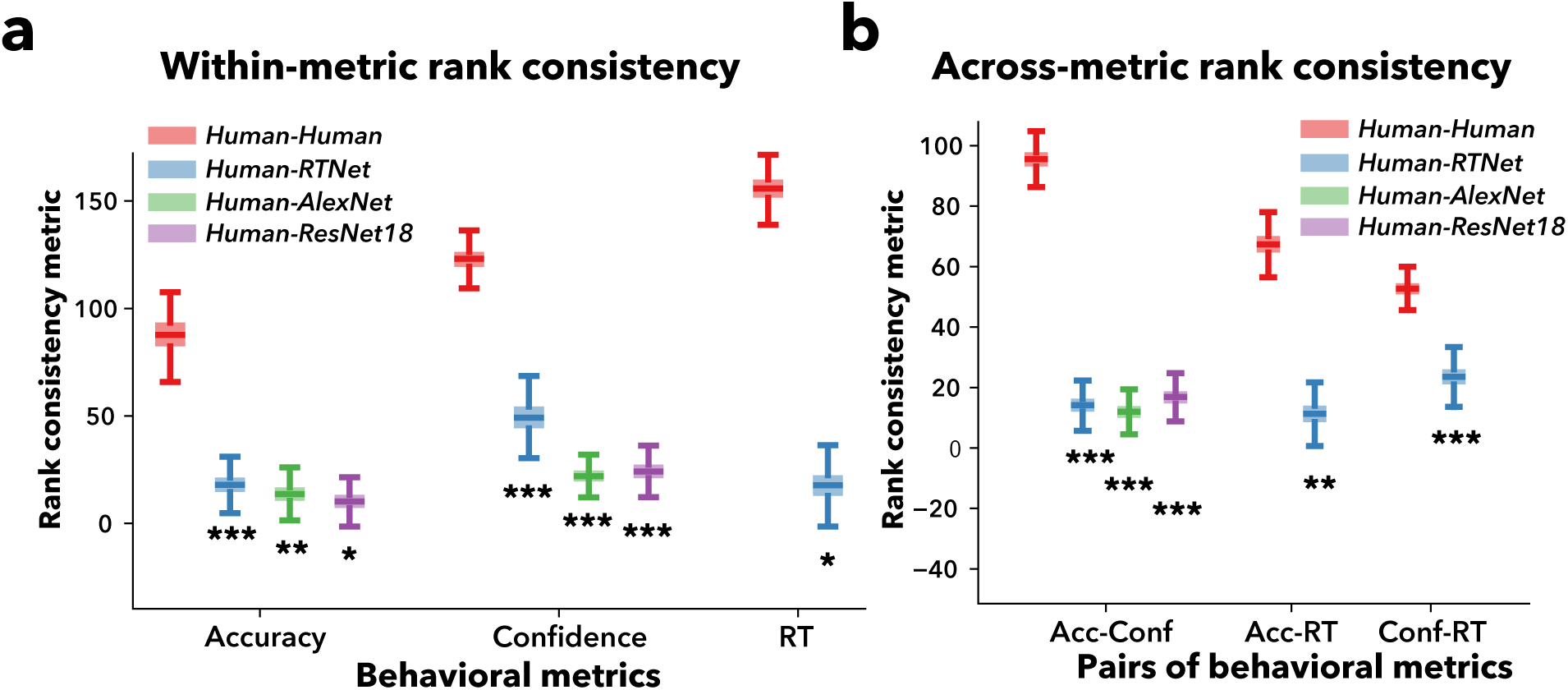
Within-metric and across-metric rank consistency results in the 10-choice object recognition task. (a) Within-metric rank consistency. (b) Across-metric rank consistency. In the main text, within-metric and across-metric consistency were quantified using correlation consistency metrics (Figure 5e,g). However, Pearson correlation may inflate consistency estimates if driven by a small number of ANN instances that consistently show poor alignment with most subjects, and this effect is not fully corrected by the subtraction. To address this concern, we computed consistency using rank ordering, based on how well each ANN instance (or each non-target subject) aligns with a target subject. These rank consistency results were largely consistent with the correlation consistency. All human-ANN mappings showed consistency significantly above chance, but lower than the human-human mapping benchmark, indicating that individual ANN instances capture some, but not all, patterns of human behavioral variability. Asterisks below the bars indicate significance against zero computed with 1000 bootstrapped samples (*, *p* < 0.05; **, *p* < 0.01; ***, *p* < 0.001).

**Supplementary Figure 10.**
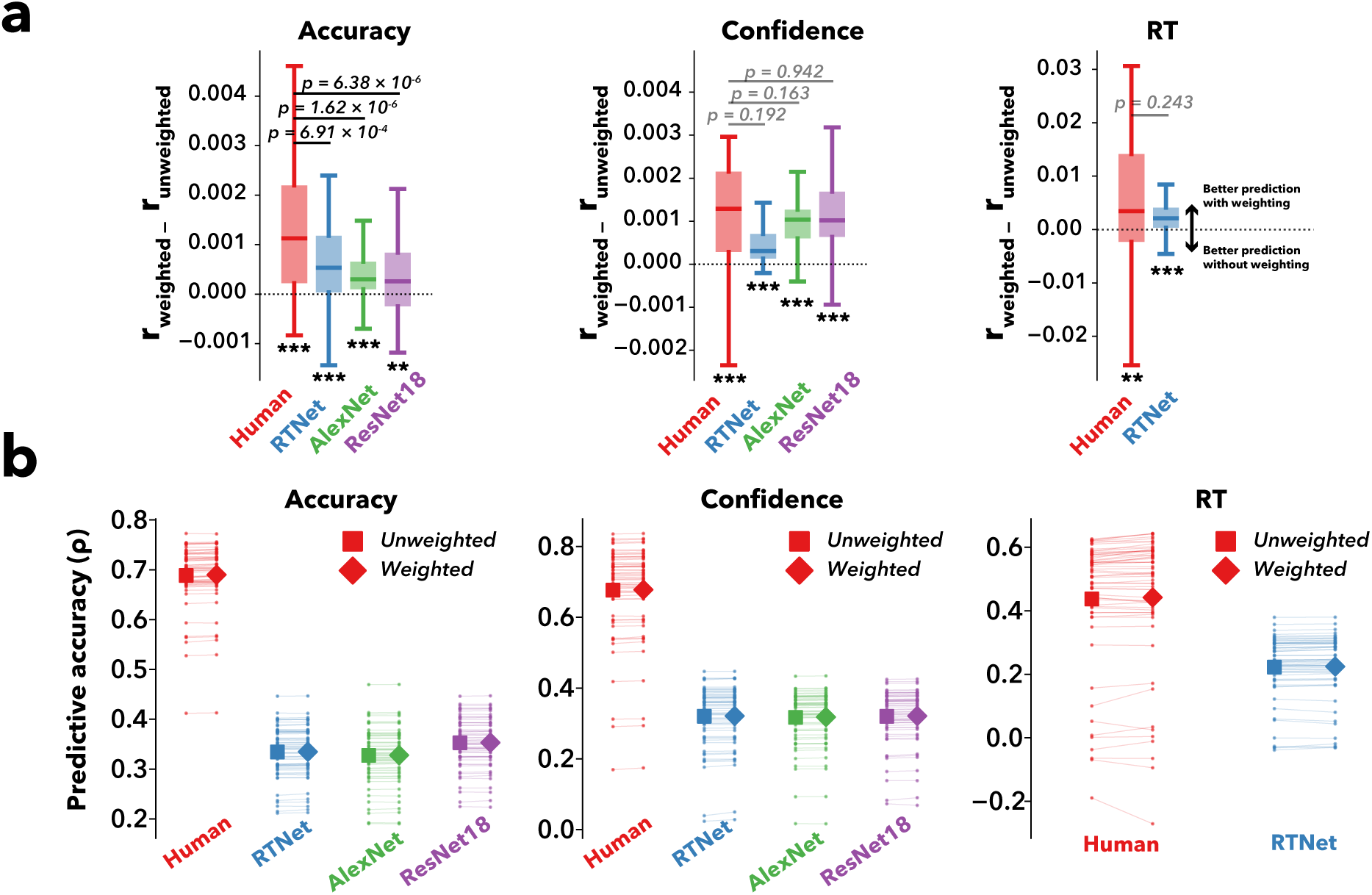
Alignment-weighted vs unweighted prediction among humans and three ANN architectures in the 10-choice object recognition task. (a) Alignment-weighted improvement. (b) Raw prediction accuracy. As in the 8-choice digit recognition task, we used the correlations in the similarity matrix as weights to predict behavior in a held-out data subset. For comparison, we computed an unweighted prediction that simply averaged across all subjects or ANN instances. All ANNs showed significant improvement in prediction with the individual alignment weighting compared to the unweighted control. We then compared these improvement from the three ANN architectures against the human benchmark. Across all three behavioral metrics, human subjects consistently exhibited higher prediction accuracy than ANNs, regardless of the weighting strategy. We found that RTNet showed a comparable level of improvement to the human benchmark for confidence and RT. Similarly, AlexNet and ResNet18 also exhibited improvement that is statistically indistinguishable from the human benchmark for confidence. However, all ANN architectures demonstrated smaller improvement than the human benchmark in accuracy. Although these improvements were modest in magnitude, they were highly consistent across different subjects. These results suggest that incorporating individual differences in ANNs could yield modest but reliable gains in predicting individual human responses to unseen data. Dotted line shows no improvement from the alignment-weighted prediction. Dots show individual human subjects. Lines connect the alignment-weighted and unweighted predictions for the same subjects. P-values indicate the significance of pairwise differences between humans and ANNs in the improvement from weighted versus unweighted predictions. Asterisks below the bars indicate significance for one-sample t-tests against zero (*, *p* < 0.05; **, *p* < 0.01; ***, *p* < 0.001; n.s., not significant).

**Supplementary Figure 11.**
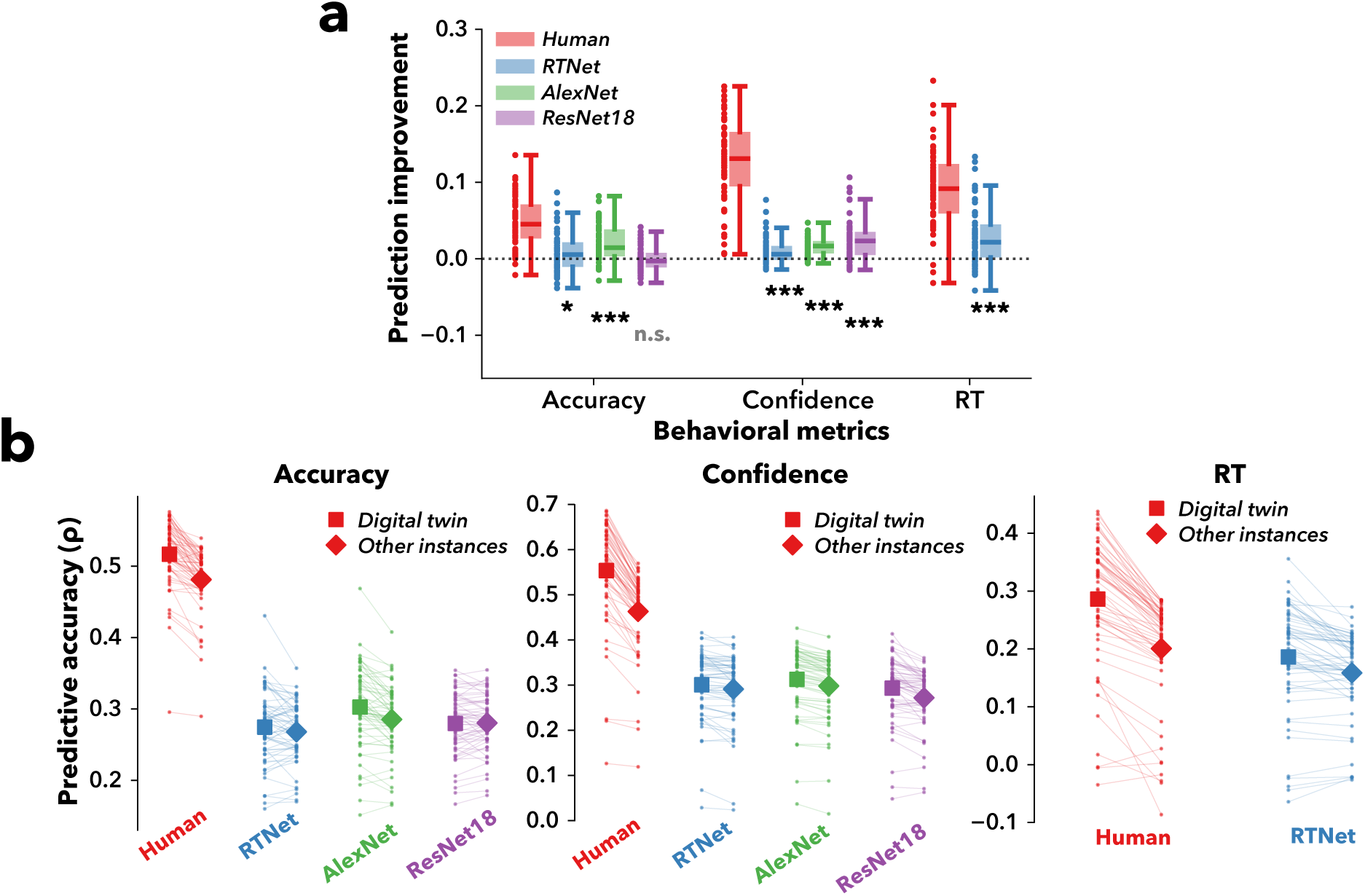
Digitial twin analysis for the 10-choice object recognition task. (a) Prediction improvement from the best-matching subject-instance pair. (b) Raw prediction accuracy for predicting single-subject responses with the digital twin vs. all other instances. As in the 8-choice digit recognition task, we identified the best-matching subject-instance pair (digital twin). We then predicted each subject’s behavior in a held-out data subset using only the responses from the digital twin. For comparison, we also predicted behavior using the responses from other instances. Human subjects showed higher prediction accuracy than ANNs for accuracy, confidence, and RT. We then analyzed the differences in improvement magnitude between the digital twin and the remaining instances. All ANNs exhibited significant improvement in predicting with digital twin except for ResNet18 in accuracy. Yet, all ANNs had their improvement magnitude lower than human subjects. Dotted line shows no improvement from the best-matching subject-instance pair. Dots show individual human subjects. Lines connect the digital twin predictions and other instances’ predictions for the same subjects. Asterisks below the bars indicate significance for one-sample t-tests against zero (*, *p* < 0.05; **, *p* < 0.01; ***, *p* < 0.001; n.s., not significant).

**Supplementary Figure 12.**
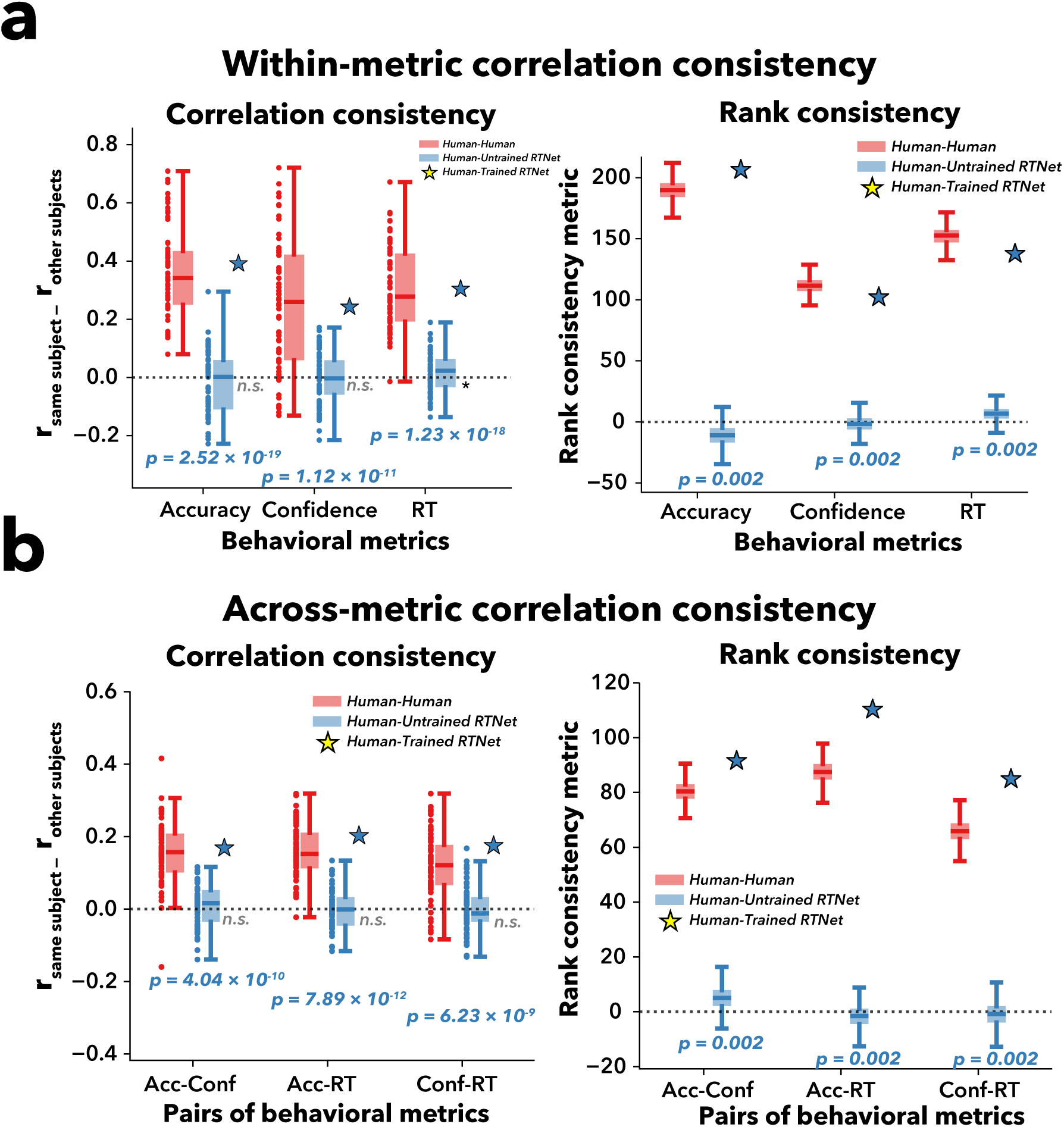
Testing individual human-ANN mapping in 60 randomly initialized untrained RTNet instances in the 8-choice digit recognition task. (a) Within-metric correlation consistency (b) Across-metric correlation consistency. To test whether network training contributes to the individual human-ANN mapping, we tested the within-metric and across-metric mapping consistency using 60 randomly initialized, untrained RTNet instances. We found that the mapping consistency of untrained RTNet instances was essentially at zero and was significantly lower compared with that of the trained RTNet instances. These results were obtained for both within-metric and across-metric mapping consistency, suggesting that network training is critical for manifesting the individual differences in neural network instances. We did not include untrained AlexNet and ResNet18 in this figure because they produced the same behavioral response for all or almost all images, preventing the computation of image-by-image correlations required for the mapping analysis. Stars indicate the average mapping consistency with trained RTNet instances that are reported in the main text (Figure 2d for within-metric & Figure 3c for across-metric). P-values indicate the statistical significance of the pairwise-comparison between the trained and untrained RTNet instances. Dotted horizontal line denotes zero consistency between two maps.

## References

Ackman, J. B., Burbridge, T. J., & Crair, M. C. (2012). Retinal waves coordinate patterned activity throughout the developing visual system. Nature, 490(7419), 219–225. 10.1038/nature11529

Annis, J., Gauthier, I., & Palmeri, T. J. (2021). Combining convolutional neural networks and cognitive models to predict novel object recognition in humans. *Journal of Experimental Psychology: Learning*, Memory, and Cognition, 47(5), 785–807. 10.1037/xlm0000968

Baek, S., Song, M., Jang, J., Kim, G., & Paik, S.-B. (2021). Face detection in untrained deep neural networks. Nature Communications, 12(1), 7328. 10.1038/s41467-021-27606-9

Bashivan, P., Kar, K., & DiCarlo, J. J. (2019). Neural population control via deep image synthesis. Science, 364(6439), eaav9436. 10.1126/science.aav9436

Bosten, J. M. (2022). Do You See What I See? Diversity in Human Color Perception. Annual Review of Vision Science, 8(Volume 8, 2022), 101–133. 10.1146/annurev-vision-093020-112820

Chow, J. K., & Palmeri, T. J. (2024). Manipulating and measuring variation in deep neural network (DNN) representations of objects. Cognition, 252, 105920. 10.1016/j.cognition.2024.105920

Conwell, C., Prince, J. S., Kay, K. N., Alvarez, G. A., & Konkle, T. (2024). A large-scale examination of inductive biases shaping high-level visual representation in brains and machines. Nature Communications, 15(1), 9383. 10.1038/s41467-024-53147-y

de Leeuw, J. R. (2015). jsPsych: A JavaScript library for creating behavioral experiments in a Web browser. Behavior Research Methods, 47(1), 1–12. 10.3758/s13428-014-0458-y

Deng, L. (2012). The MNIST Database of Handwritten Digit Images for Machine Learning Research [Best of the Web]. IEEE Signal Processing Magazine, 29(6), 141–142. 10.1109/MSP.2012.2211477

Feather, J., Khosla, M., Murty, N. A. R., & Nayebi, A. (2025). Brain-Model Evaluations Need the NeuroAI Turing Test (arXiv:2502.16238). arXiv. 10.48550/arXiv.2502.16238

Fung, H., Shekhar, M., Xue, K., Rausch, M., & Rahnev, D. (2025). Similarities and differences in the effects of different stimulus manipulations on accuracy and confidence. Consciousness and Cognition, 136, 103942. 10.1016/j.concog.2025.103942

Gao, W., Elton, A., Zhu, H., Alcauter, S., Smith, J. K., Gilmore, J. H., & Lin, W. (2014). Intersubject variability of and genetic effects on the brain’s functional connectivity during infancy. Journal of Neuroscience, 34(34), 11288–11296. https://www.jneurosci.org/content/34/34/11288.short

Green, M. L., Hu, M., Denison, R. N., & Rahnev, D. (2026). Using Artificial Neural Networks to Relate External Sensory Features to Internal Decisional Evidence. Open Mind, 10, 29–46. 10.1162/OPMI.a.317

Hanganu, I. L., Ben-Ari, Y., & Khazipov, R. (2006). Retinal Waves Trigger Spindle Bursts in the Neonatal Rat Visual Cortex. The Journal of Neuroscience, 26(25), 6728–6736. 10.1523/JNEUROSCI.0752-06.2006

He, K., Zhang, X., Ren, S., & Sun, J. (2015). Deep Residual Learning for Image Recognition (arXiv:1512.03385). arXiv. 10.48550/arXiv.1512.03385

Jang, H., & Tong, F. (2024). Improved modeling of human vision by incorporating robustness to blur in convolutional neural networks. Nature Communications, 15(1), 1989. 10.1038/s41467-024-45679-0

Jinsi, O., Henderson, M. M., & Tarr, M. J. (2023). Early experience with low-pass filtered images facilitates visual category learning in a neural network model. PLOS ONE, 18(1), e0280145. 10.1371/journal.pone.0280145

Jordan, K. (2024). On the Variance of Neural Network Training with respect to Test Sets and Distributions (arXiv:2304.01910). arXiv. 10.48550/arXiv.2304.01910

Kriegeskorte, N. (2015). Deep Neural Networks: A New Framework for Modeling Biological Vision and Brain Information Processing. Annual Review of Vision Science, 1(1), 417–446. 10.1146/annurev-vision-082114-035447

Krizhevsky, A., Sutskever, I., & Hinton, G. E. (2012). Imagenet classification with deep convolutional neural networks. Advances in Neural Information Processing Systems, 25. https://proceedings.neurips.cc/paper/2012/hash/c399862d3b9d6b76c8436e924a68c45b-Abstract.html

Li, Q., Joo, S. J., Yeatman, J. D., & Reinecke, K. (2020). Controlling for Participants’ Viewing Distance in Large-Scale, Psychophysical Online Experiments Using a Virtual Chinrest. Scientific Reports, 10(1), Article 1. 10.1038/s41598-019-57204-1

Linsley, D., Feng, P., & Serre, T. (2025). Better artificial intelligence does not mean better models of biology. Trends in Cognitive Sciences, 0(0). 10.1016/j.tics.2025.11.016

Lu, Z., Thorat, S., Cichy, R. M., & Kietzmann, T. C. (2025). Adopting a human developmental visual diet yields robust*, shape-based AI vision* (arXiv:2507.03168). arXiv. 10.48550/arXiv.2507.03168

McGrath, S. W., Russin, J., Pavlick, E., & Feiman, R. (2024). How Can Deep Neural Networks Inform Theory in Psychological Science? Current Directions in Psychological Science.

McNeal, N., Deb, M., & Murty, A. R. (2024, October 10). Small-scale adversarial perturbations expose differences between predictive encoding models of human fMRI responses. UniReps: 2nd Edition of the Workshop on Unifying Representations in Neural Models. https://openreview.net/forum?id=1g7HWFmvY0#discussion

Mehrer, J., Spoerer, C. J., Jones, E. C., Kriegeskorte, N., & Kietzmann, T. C. (2021). An ecologically motivated image dataset for deep learning yields better models of human vision. Proceedings of the National Academy of Sciences, 118(8), e2011417118. 10.1073/pnas.2011417118

Mehrer, J., Spoerer, C. J., Kriegeskorte, N., & Kietzmann, T. C. (2020). Individual differences among deep neural network models. Nature Communications, 11(1), 5725. https://www.nature.com/articles/s41467-020-19632-w

Mueller, K. N., Carter, M. C., Kansupada, J. A., & Ponce, C. R. (2023). Macaques recognize features in synthetic images derived from ventral stream neurons. Proceedings of the National Academy of Sciences, 120(10), e2213034120. 10.1073/pnas.2213034120

Ponce, C. R., Xiao, W., Schade, P. F., Hartmann, T. S., Kreiman, G., & Livingstone, M. S. (2019). Evolving Images for Visual Neurons Using a Deep Generative Network Reveals Coding Principles and Neuronal Preferences. Cell, 177(4), 999–1009.e10. 10.1016/j.cell.2019.04.005

Rafiei, F., Shekhar, M., & Rahnev, D. (2024). The neural network RTNet exhibits the signatures of human perceptual decision-making. Nature Human Behaviour. 10.1038/s41562-024-01914-8

Rahnev, D. (2021). Response Bias Reflects Individual Differences in Sensory Encoding. Psychological Science, 32(7), 1157–1168. 10.1177/0956797621994214

Ratan Murty, N. A., Bashivan, P., Abate, A., DiCarlo, J. J., & Kanwisher, N. (2021). Computational models of category-selective brain regions enable high-throughput tests of selectivity. Nature Communications, 12(1), 5540. 10.1038/s41467-021-25409-6

Richler, J. J., Tomarken, A. J., Sunday, M. A., Vickery, T. J., Ryan, K. F., Floyd, R. J., Sheinberg, D., Wong, A. C.-N., & Gauthier, I. (2019). Individual differences in object recognition. Psychological Review, 126(2), 226. https://psycnet.apa.org/record/2019-09750-002

Shahbazi, E., Ma, T., Pernuš, M., Scheirer, W., & Afraz, A. (2024). Perceptography unveils the causal contribution of inferior temporal cortex to visual perception. Nature Communications, 15(1), 3347. 10.1038/s41467-024-47356-8

Shekhar, M., Fung, H., Saxena, K., Rafiei, F., & Rahnev, D. (2025). Using artificial neural networks to reveal the human confidence computation. PLOS Computational Biology, 21(12), e1013827. 10.1371/journal.pcbi.1013827

Shekhar, M., & Rahnev, D. (2024). Human-like dissociations between confidence and accuracy in convolutional neural networks. PLOS Computational Biology, 20(11), e1012578. 10.1371/journal.pcbi.1012578

Strock, A., Mistry, P. K., & Menon, V. (2025). Personalized deep neural networks reveal mechanisms of math learning disabilities in children. Science Advances, 11(23), eadq9990. 10.1126/sciadv.adq9990

Turing, A. M. (1950). Computing machinery and intelligence. Mind, 59, 433–460. https://www.edwardfrenkel.com/turing-intelligence.pdf

Vogelsang, M., Vogelsang, L., Gupta, P., Gandhi, T. K., Shah, P., Swami, P., Gilad-Gutnick, S., Ben-Ami, S., Diamond, S., Ganesh, S., & Sinha, P. (2024). Impact of early visual experience on later usage of color cues. Science, 384(6698), 907–912. 10.1126/science.adk9587

Walker, E. Y., Sinz, F. H., Cobos, E., Muhammad, T., Froudarakis, E., Fahey, P. G., Ecker, A. S., Reimer, J., Pitkow, X., & Tolias, A. S. (2019). Inception loops discover what excites neurons most using deep predictive models. Nature Neuroscience, 22(12), 2060–2065. 10.1038/s41593-019-0517-x

Wichmann, F. A., & Geirhos, R. (2023). Are Deep Neural Networks Adequate Behavioral Models of Human Visual Perception? Annual Review of Vision Science, 9(1), 501–524. 10.1146/annurev-vision-120522-031739

Wong, R. O. L. (1999). Retinal waves and visual system development. Annual Review of Neuroscience, 22(Volume 22, 1999), 29–47. 10.1146/annurev.neuro.22.1.29

Wood, J. N., Pandey, L., & Wood, S. M. W. (2024). Digital Twin Studies for Reverse Engineering the Origins of Visual Intelligence. Annual Review of Vision Science, 10(Volume 10, 2024), 145–170. 10.1146/annurev-vision-101322-103628

Xu, Y., Cao, M., Liao, X., Xia, M., Wang, X., Jeon, T., Ouyang, M., Chalak, L., Rollins, N., Huang, H., & He, Y. (2019). Development and Emergence of Individual Variability in the Functional Connectivity Architecture of the Preterm Human Brain. Cerebral Cortex, 29(10), 4208–4222. 10.1093/cercor/bhy302

